# Optimizing genomic prediction for complex traits via investigating multiple factors in switchgrass

**DOI:** 10.1101/2024.06.17.599440

**Authors:** Peipei Wang, Fanrui Meng, Christina B Del Azodi, Kenia Segura Abá, Michael D. Casler, Shin-Han Shiu

## Abstract

Genomic prediction, the use of genetic information for predicting traits, has accelerated the breeding processes and provided mechanistic insights into the genetic bases of complex traits. While substantial efforts have been devoted to optimize genomic prediction, there remain areas that need to be further explored, including the impact of genome assemblies, genotyping approaches, variant types, allelic complexities, polyploidy levels, and population structures. Here, we assess the impact of these factors on the prediction of 20 complex traits in switchgrass (*Panicum virgatum* L.), a perennial biofuel feedstock. We found that short read-based genome assembly perform comparably to or even better than long read-based assembly in trait prediction; exome capture-based models have higher prediction accuracy than genotyping-by-sequencing-based models for 13 traits; bi-allelic insertion/deletions are as useful as bi-allelic single nucleotide polymorphisms in trait prediction, whereas multi-allelic variants outperform bi-allelic ones for 15 traits. Models built for tetraploids have higher prediction accuracy than those for octoploids for most traits. Traits of individuals with higher within-population genetic distances tend to have higher prediction accuracy. Finally, integrating different types of variants can improve the prediction accuracy. By exploring these factors, anthesis date prediction models built using multi-allelic insertion/deletions derived from exome capture led to the largest number of orthologs of benchmark flowering time genes compared to other models. Our study provides insights into the factors influencing genomic prediction outcomes that inform best practices for future studies and for improving agronomic traits in switchgrass and other species through selective breeding.

## Introduction

Before the genomic era, the breeding process relied heavily on natural variation, and hybridization or mutagenesis, and was time- and labor-consuming. In the genomic era, the molecular variant-assisted breeding technologies, especially genomic selection or genomic prediction (GP), have dramatically accelerated the breeding processes for both animals and plants (Crossa et al., 2017; Alemu et al., 2024; Tade and Melesse, 2024). Learning from a training population with both genetic variants (e.g., single nucleotide polymorphisms, SNPs) and phenotypic data (e.g., flowering time), GP establishes statistical models to build the connection between the genetic variants and phenotypic data, which can be applied to a new population that has been genotyped but with unmeasured phenotypes. This can be done without planting all the seedlings and waiting for a whole growing season, therefore resulting in shortened breeding cycles (Estopa et al., 2023; Alemu et al., 2024).

Substantial effort has been made to understand the effects of different factors on the accuracy of GP (Norman et al., 2018; Zhang et al., 2019; Kriaridou et al., 2020; Batista et al., 2022; Alemu et al., 2024; Zheng et al., 2024), including variant density (Kriaridou et al., 2020), population size (Akdemir et al., 2019; Wu et al., 2023), population structure (Norman et al., 2018), relationship between training and test populations (Isidro et al., 2015), functional genomic annotations (Zheng et al., 2024), marker ploidy (Aalborg and Nielsen, 2024), variants besides SNPs (e.g., structural variants, Liu et al., 2024) and statistical methods (Azodi et al., 2019; Zhang et al., 2019; Wang et al., 2023b; Alemu et al., 2024), etc. For example, the prediction accuracy increases along with the increase of variant density to a certain point (Norman et al., 2018; Kriaridou et al., 2020). Regarding the statistical methods used in GP, a relatively simple linear model, ridge regression Best Linear Unbiased Prediction (rrBLUP), performed similarly or better compared with other “state-of-the-art” approaches for most scenarios, such as random forest, support vector machine, gradient boosting, artificial neural net, and convolutional neural net (Azodi et al., 2019; Azodi et al., 2020; Wang et al., 2024). Beyond the above, there are additional factors with their effects on GP accuracy remaining largely unexplored, such as genome assemblies, genotyping approaches, variant allelic complexities, and so on. With reduced sequencing cost and improvement in assembly algorithms, a number of plant species have genome assemblies improved from scaffold-level to chromosome-level or even telomere-to-telomere (T2T) (Chen et al., 2023; Lu et al., 2024). However, the higher the quality of the assembly, the more the cost needed, and it remains unknown whether a better assembly is necessarily to improve the GP accuracy since few studies have focused on the effects of assembly quality on GP accuracy. A blueberry study showed that the chromosome-level assembly ‘Draper’ led to comparable prediction accuracy for most traits as the scaffold-level assembly ‘W8520’ with only half numbers of probes (Benevenuto et al., 2019), but it was still unknown whether the chromosome-level assembly led to improved GP accuracy or not when the genome-wide variants were used.

A number of approaches have been developed for genotyping, including SNP array, genotyping by random amplicon sequencing-direct system, genotyping-by-sequencing, exome capture, whole genome sequencing (WGS), and others (Scheben et al., 2017; Ayalew et al., 2022; Boatwright et al., 2022; Wang et al., 2023a; Minamikawa et al., 2024). WGS covers the majority of genetic variants across the whole genome, whereas the other approaches only cover a portion of variants but with relatively lower costs. It remains unclear how different genotyping approaches impact the GP accuracy and which approaches would be a better option with different aims of the studies.

The genetic variants can be both bi-allelic and multi-allelic: out of the SNPs called for the Arabidopsis 1001G data using the 250k SNP-array from Kim et al. (2007), 7% were multi-allelic (Alonso-Blanco et al., 2016); out of the SVs identified using 32 *A. thaliana* ecotypes, multi-allelic SVs (29.65 Mb) were much more than bi-allelic SVs (7.51 Mb, Kang et al., 2023). However, the majority of past genome-wide association study (GWAS) and GP studies mainly used bi-allelic variants (Crossa et al., 2017; Misra et al., 2017; Song et al., 2018; Wang et al., 2023b). It is important to know how the multi-allelic variants contribute to complex trait prediction to make the best of the ever increasing genomic data, such as graph-based pan-genomes that are used as references to call all types of genetic variants with high quality.

Here, using switchgrass (*Panicum virgatum* L.)—a perennial grass with both tetra- and octo-ploids and a key species as a bioenergy feedstock (McLaughlin and Adams Kszos, 2005)—as a model system, we aim to optimize GP by assessing the impacts on GP accuracy due to differences in: 1) genome assembly versions, 2) genotyping strategies, 3) genetic variant types, 4) numbers of variant alleles, and 5) polyploids and population structures. Both short-read and long-read based assemblies are available for switchgrass, and there are two genotyping datasets for a switchgrass diversity panel with 486 individuals: GBS (Lu et al., 2013) and EC (Evans et al., 2015). We also assess how different types of genetic variants impact our ability to uncover the genetic basis underlying anthesis date by interpreting GP models.

## Results

### Baseline models predicted trait values with variable accuracy

There are 20 phenotypic trait values available for the switchgrass diversity panel we used (**Table S1**), including 7 morphological and 13 biochemical traits (**Table S2**, Lipka *et al*., 2014). Among the 20 traits, some are highly associated with each other, such as three biochemical traits, namely cell wall concentration, etherified ferulates and *in vitro* dry matter digestibility, and two flowering time-related traits, heading date and anthesis date (**Fig. 1A**). To determine how well these 20 phenotypic traits can be predicted in switchgrass, we first built GP models using GBS bi-allelic SNPs that were mapped to the version 5 genome assembly of the lowland tetraploid AP13 (Lovell et al., 2021). The rrBLUP approach was used to build the models due to its relatively better or similar prediction accuracy compared with other methods, and its interpretability for underlying molecular mechanisms for target traits (Azodi et al., 2019; Azodi et al., 2020; Wang et al., 2024). Information of 20% of the individuals in the diversity panel were held out as the test set before model building to serve as an independent dataset for evaluating the performance of final models. The data of the remaining 80% individuals were used to train the models with a five-fold cross-validation scheme (see **Methods**). To compare prediction accuracy among different traits, we also established models using the population structure (p; defined as the first five principal components of the corresponding genetic variants [g], see **Methods**) to determine the baseline prediction accuracy for each trait (Azodi et al., 2020). The r^2^ of the Pearson Correlation Coefficient (PCC) between the true and predicted trait values for individuals were first calculated for models built using genetic variants (r^2^_g_) and population structure (r^2^_p_), separately. Then the improvement in r^2^ (r^2^_i_ = r^2^_g_ - r^2^_p_) was used to measure the performance of the GP models. This r^2^_i_ was calculated both for individuals in the cross-validation (CV) set (r^2^_i,CV_) and the test set (r^2^_i,test_).

**Fig. 1.**
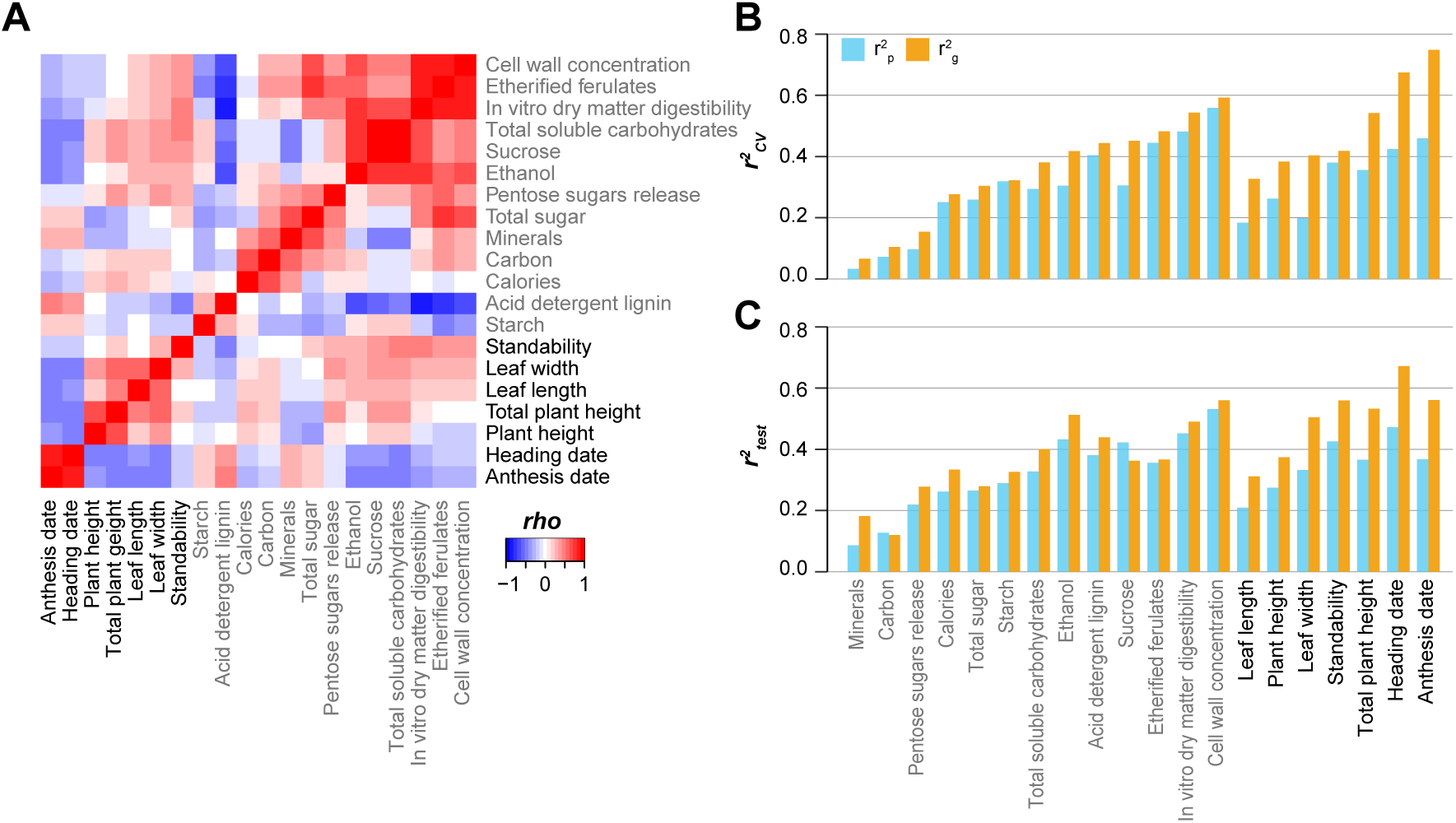
Correlation between the values of 20 traits and the prediction accuracy for these 20 traits. (**A**) Heatmap showing the correlation between the values of 20 traits. Gray font: 13 biochemical traits; black font: seven morphological traits. Color scale: rho of the spearman’s rank correlation coefficient. (**B**,**C**) Prediction accuracy (r^2^) for 20 traits in cross-validation (**B**) and test (**C**) subsets for models built using the bi-allelic single nucleotide polymorphisms (SNPs) called by mapping the genotyping-by-sequencing (GBS) data to v5 genome assembly. The models were built on all individuals including tetraploids and octoploids. X-axis: 20 traits; y-axis: median r^2^ between true and predicted trait values among 10 replicate runs. Blue (r^2^_p_): r^2^ for models built with the population structure, which was defined as the first five principal components from the genetic variants; orange (r^2^_g_): r^2^ of models built with genetic variants.

We found that r^2^_i,CV_ was positively correlated with r^2^_i,test_ (PCC=0.68, *p*-value=9.89e-04, **Fig. 1B,C**), suggesting the relative robustness of our models. In general, r^2^_i,CV_ for morphological traits (average r^2^_i,CV_=0.176) were higher than those for biochemical traits (average r^2^_i,CV_=0.055, *p*-value of Wilcoxon signed rank test=4.66e-03), and the two flowering time related traits—anthesis date and heading date—had the highest r^2^_i,CV_ (0.289 and 0.250, respectively). These results indicate that morphological traits tended to have higher heritability, whereas the prediction accuracy (r^2^_g_) for most biochemical traits were mainly confounded by population structure. Thus, cautions need to be taken in the GWAS and GP practices for those biochemical traits.

### Better genome assembly did not provide better trait prediction

To test whether the genome assembly with improved quality (i.e., v5, Lovell et al., 2021) would lead to higher prediction accuracy for switchgrass traits than short read-based assembly (e.g., v1), we also mapped GBS sequences to the v1 assembly and called the genetic variants using the same methods and parameter settings as those for v5-based variants (see **Methods**). The assembly sizes of v1 (1,230 Mb) and v5 assemblies (1,129 Mb) were similar, and there were almost equivalent numbers of variants for v1 (11,042) and v5 (11,020) assemblies, with similar distribution of variants across different gene functional regions (**Fig. S1A,B** and **Table S3**). In addition, most of the GBS bi-allelic SNPs were shared between these two assemblies, as 5,283 (69.92%) out of 7,556 v1-based bi-allelic SNPs had corresponding v5-based ones (see **Methods** and **Table S4**). Beyond these similarities, N50 for v5 assembly (N50=5.5 Mb) is nearly 100 times higher than v1 (N50=54 Kb). However, v5-based models outperformed v1-based models only for anthesis date, with the average difference in r^2^_i,CV_ between two models for 20 traits was only 0.006 (the “All” column in **Fig. 2A**, **Fig. 2B**; for the r^2^_p,CV_, r^2^_g,CV_, r^2^_i,CV_, r^2^_p,test_, r^2^_g,test_, and the r^2^_i,test_, see **Fig. S2-7**, respectively). This result suggests that the improvement in contig size may not be relevant to GP because genetic variants are treated as independent variables in GP algorithms.

**Fig. 2.**
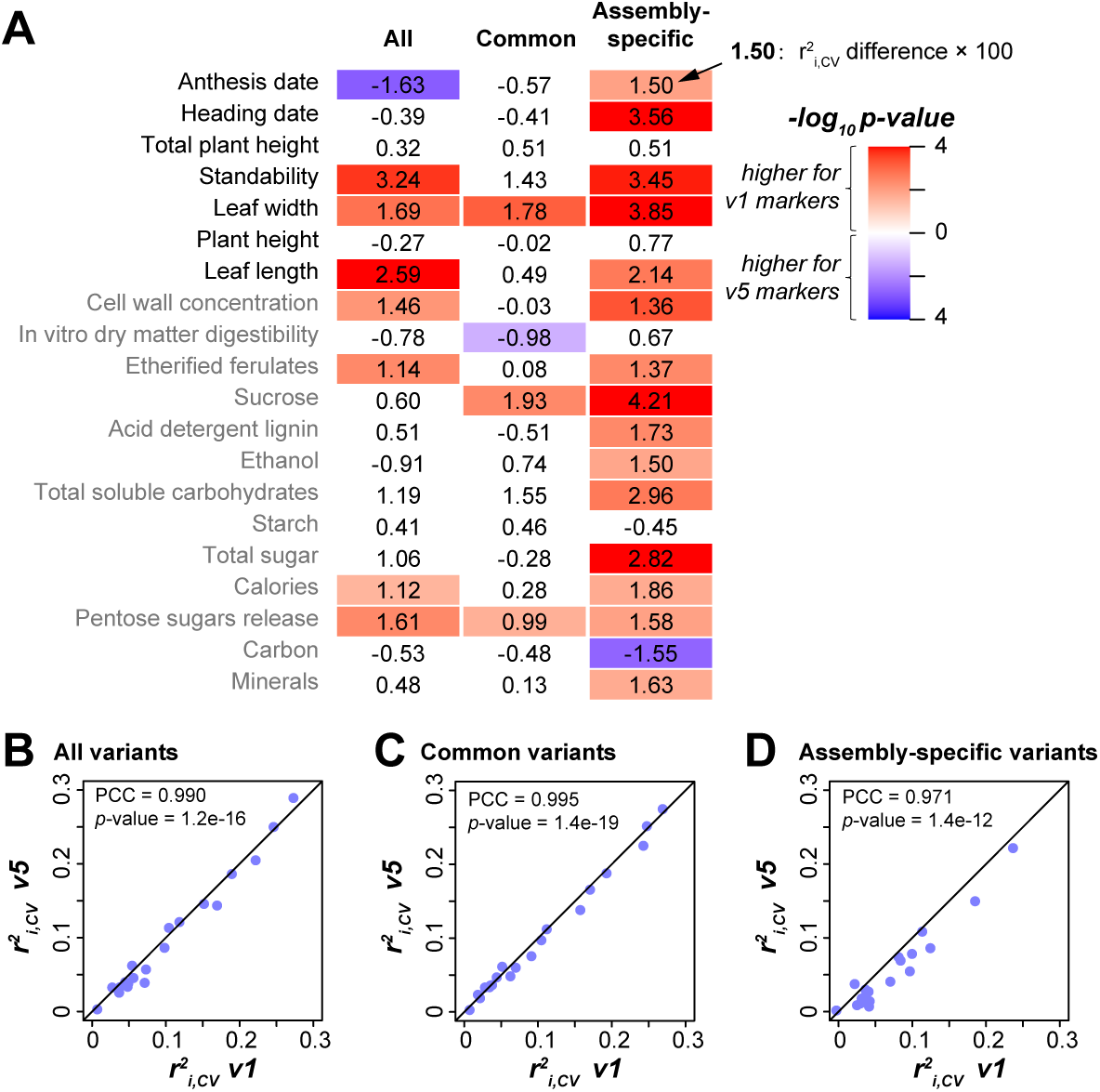
Prediction accuracy differences between models based on v1 and v5 assemblies. (**A**) Differences in r^2^_i,CV_ between v1- and v5-based models. The column “All”: models built using all the v1- and v5-based GBS bi-allelic SNPs; “Common”: models built using the common GBS bi- allelic SNPs shared between v1 and v5 assemblies; “Assembly-specific”: models built using GBS bi-allelic SNPs that are specific to v1 and v5 assemblies, separately. Values in the heatmap: r^2^_i,CV_ differences × 100; only the comparisons with statistical significance (*p* from Wilcoxon signed-rank test < 0.05) are indicated with colors; color scale: -log_10_ *p*-value; red and blue: significantly higher r^2^_i,CV_ was observed for the v1- and v5-based models, respectively. (**B**-**D**) Correlation between r^2^_i,CV_ of v1-(x-axis) and v5-based (y-axis) models. (**B**) Models built using all the v1- and v5-based GBS bi-allelic SNPs; (**C**) Models built using the common GBS bi-allelic SNPs shared between v1 and v5 assemblies; (**D**) Models built using GBS bi-allelic SNPs that are specific to v1 and v5 assemblies, separately.

In fact, although the effect sizes (differences in r^2^_i,CV_) were small, models built with variants derived from v1 had significantly better r^2^_i,CV_ than v5-based models for seven traits (the “All” column in **Fig. 2A**). These differences in r^2^_i,CV_ can be partially explained by the assembly-specific variants: models built using common variants (shared between v1 and v5 assemblies) had similar performance (the “Common” column in **Fig. 2A**, **Fig. 2C**), while assembly-specific variants-based models tended to have better performance when they were v1-based (the “Assembly-specific” column in **Fig. 2A**, **Fig. 2D**). A possible explanation of this finding that needs to be investigated further is that, while v5 assembly had improved contig sizes, some genomic regions present in the v1 assembly were not assembled from long-reads and thus were no longer present in the v5 assembly. These results suggest that, in switchgrass, short-read-based assembly is equally competent as or even superior to long-read-based assembly for the purpose of GP. Nevertheless, since the v5 assembly is used by the community, our subsequent analyses are based on genetic variants called using the v5 assembly.

### Exome capture SNPs led to models better than those based on GBS

EC and GBS are two commonly used genotyping methods to capture the genetic variants with a much lower cost than WGS, but use different strategies: EC identifies genetic variants that may alter protein sequences, while GBS captures variants around restriction sites of (a) given restriction enzyme(s). We found that when using bi-allelic SNPs, EC-based models had significantly higher r^2^_i,CV_ than GBS-based models for 15 traits but lower r^2^_i,CV_ for only two traits (the “Unbalanced” column, **Fig. 3A**). Considering that there were ∼72 times more EC variants (526,705, **Fig. S1C**) than GBS ones (7,357), to eliminate the influence of variant numbers on prediction accuracy, we randomly down-sampled EC bi-allelic SNPs to the same number of GBS bi-allelic SNPs, and conducted this down-sampling 100 times (**Methods**). The median r^2^_i,CV_ of the 100 models built with these down-sampled EC variants still had significantly higher r^2^_i,CV_ than GBS-based models for 13 traits, but had significantly lower r^2^_i,CV_ than GBS-based models for only three traits (the “Balanced” column, **Fig. 3A**). These results suggest that, regardless of the numbers of variants, the distribution of variants may be the main factor influencing GP accuracy when comparing EC and GBS-based models.

**Fig. 3.**
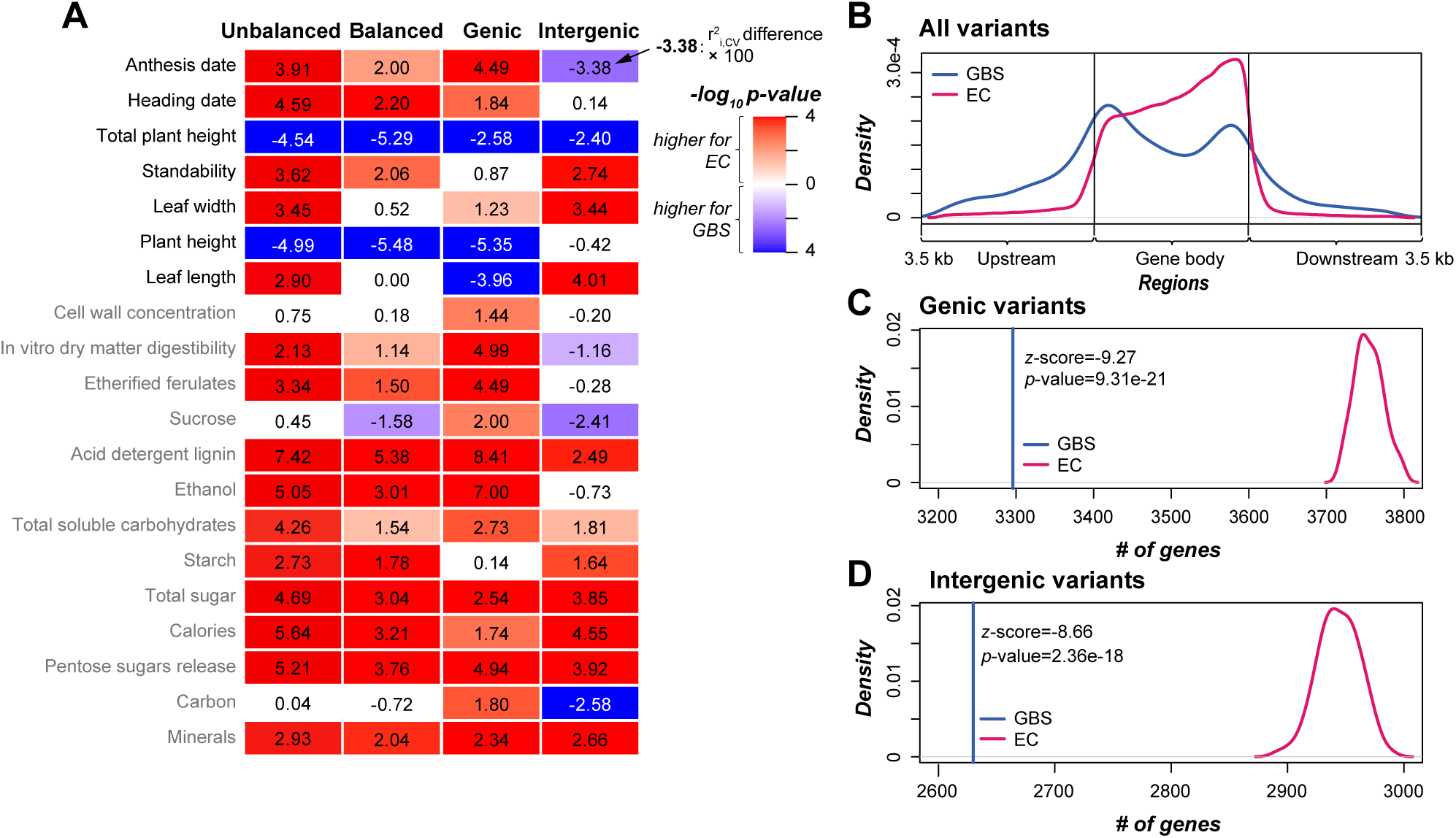
Better trait prediction in exome capture (EC)-based models than genotyping-by- sequencing (GBS)-based models. (**A**) Differences in r^2^_i,CV_ between EC- and GBS-based models. The column “Unbalanced”: models built using all EC and GBS bi-allelic SNPs; “Balanced”: models built using balanced EC (down-sampled to match the number of GBS variants) and GBS bi-allelic SNPs; “Genic”: models built using balanced EC and GBS bi-allelic SNPs that were located within the gene bodies; “Intergenic”: models built using balanced EC and GBS bi-allelic SNPs that were located within intergenic regions. Values in the heatmap: r^2^_i,CV_ differences × 100; only the comparisons with statistical significance (*p* from Wilcoxon signed-rank test < 0.05) are indicated with colors; color scale: -log_10_ *p*-value; red and blue: significantly higher r^2^_i,CV_ was observed for the EC- and GBS-based models, respectively. (**B**) Distribution of the GBS (blue) and EC (pink) bi-allelic SNPs in the genome. All the gene bodies were scaled to 3.5 Kb to make the locations of variants within the gene body comparable. The regions of 3.5 Kb up-stream and down-stream the gene bodies were also shown. (**C,D**) Comparison between the numbers of genes harboring (**C**) or adjacent to (**D**) GBS bi-allelic SNPs (blue line) and the downsampled EC bi-allelic SNPs (pink line). z-score values of the GBS-based numbers in the distribution of EC- based numbers are shown in the figures.

We found that EC variants tended to be located in genic regions or in intergenic regions that are closer to genes (**Fig. 3B, Fig. S7**), and contained more genic variants (86.4%) than GBS variants (55.5%, **Table S3**). We further established GP models using only genic GBS and EC variants (down-sampled to match the number of genic GBS variants) and found that genic GBS- based models still had significantly lower r^2^_i,CV_ than genic EC-based models for 15 of 20 traits (the “Genic” column, **Fig. 3A**). When examining the GBS bi-allelic SNPs and the 100 down-sampled subsets of EC bi-allelic SNPs in detail, we found significantly fewer genes that contained GBS variants (3,296) than contained EC variants (median across 100 replicates=3,754; *p*- value=9.31e-21, **Fig. 3C**). These findings indicate that having higher gene coverage, rather than more genic variants, may explain the higher GP accuracy of EC-based models.

Furthermore, we also built models using intergenic GBS and EC bi-allelic SNPs, and found that GBS-based models had significantly smaller r^2^_i,CV_ than EC-based models (balanced for variant numbers) for 10 of 20 traits, but significantly higher r^2^_i,CV_ only for five traits (the “Intergenic” column, **Fig. 3A**). Similar to genic variants, there were fewer genes adjacent to (3.5 Kb up- or downstream the variants) intergenic GBS variants (2,630, *p*-value=2.36e-18, **Fig. 3D**) than intergenic EC variants (down-sampled to match intergenic GBS, the median number of genes among 100 replicates was 2,944). This result suggests that, beyond higher gene coverage, including variants that are adjacent to more genes may also improve the GP accuracy, which is potential due to the linkage between these intergenic variants and genes. Another potential explanation for the better performance of models built using intergenic EC variants was because the intergenic EC variants were closer to gene bodies than the intergenic GBS variants (**Fig. 3B**). If this was true, we would expect negative correlations between the distance to the gene body and the absolute coefficient of variants that was used as a proxy for the contribution of variants to trait prediction. However, we did not see such correlation for leaf width models as an example (Spearman’s rank correlation coefficients = -7.7e-03 and 7.2e-03 for models built with intergenic GBS and EC bi-allelic SNPs, respectively), for which the EC-based model had significantly higher r^2^_i,CV_ than the GBS-based model (the “Intergenic” column, **Fig. 3A**). Taken together, those results suggest that EC variants are superior to GBS variants in predicting switchgrass traits, and gene coverage by variants has a significant impact on how well a trait can be predicted.

### Multi-allelic variants outperformed bi-allelic variants in trait prediction

Up to this point, all the genetic variants we have explored are bi-allelic SNPs, which are commonly used in GWAS and GP studies. Besides bi-allelic SNPs, bi-allelic indels and multi- allelic variants also contribute to phenotypic variation, and are informative for trait prediction (Veerkamp et al., 2016; Biová et al., 2024), but with the degree to which they contribute to GP accuracy largely unexplored. For the GBS data, compared with the 7,357 bi-allelic SNPs, 1,827 bi-allelic indels were identified (**Table S3**). Models built using bi-allelic indels had significantly lower r^2^_i,CV_ than SNP-based models for 15 traits (without down-sampling, the 1^st^ column, **Fig. 4A)**, but had comparable performance as balanced SNP-based models (with lower r^2^_i,CV_ for 6 traits and higher r^2^_i,CV_ for another 6 traits, the 2^nd^ column, **Fig. 4A**). These results suggest that, although occur less frequently than SNPs, indels are comparably useful as SNPs when the numbers are balanced.

**Fig. 4.**
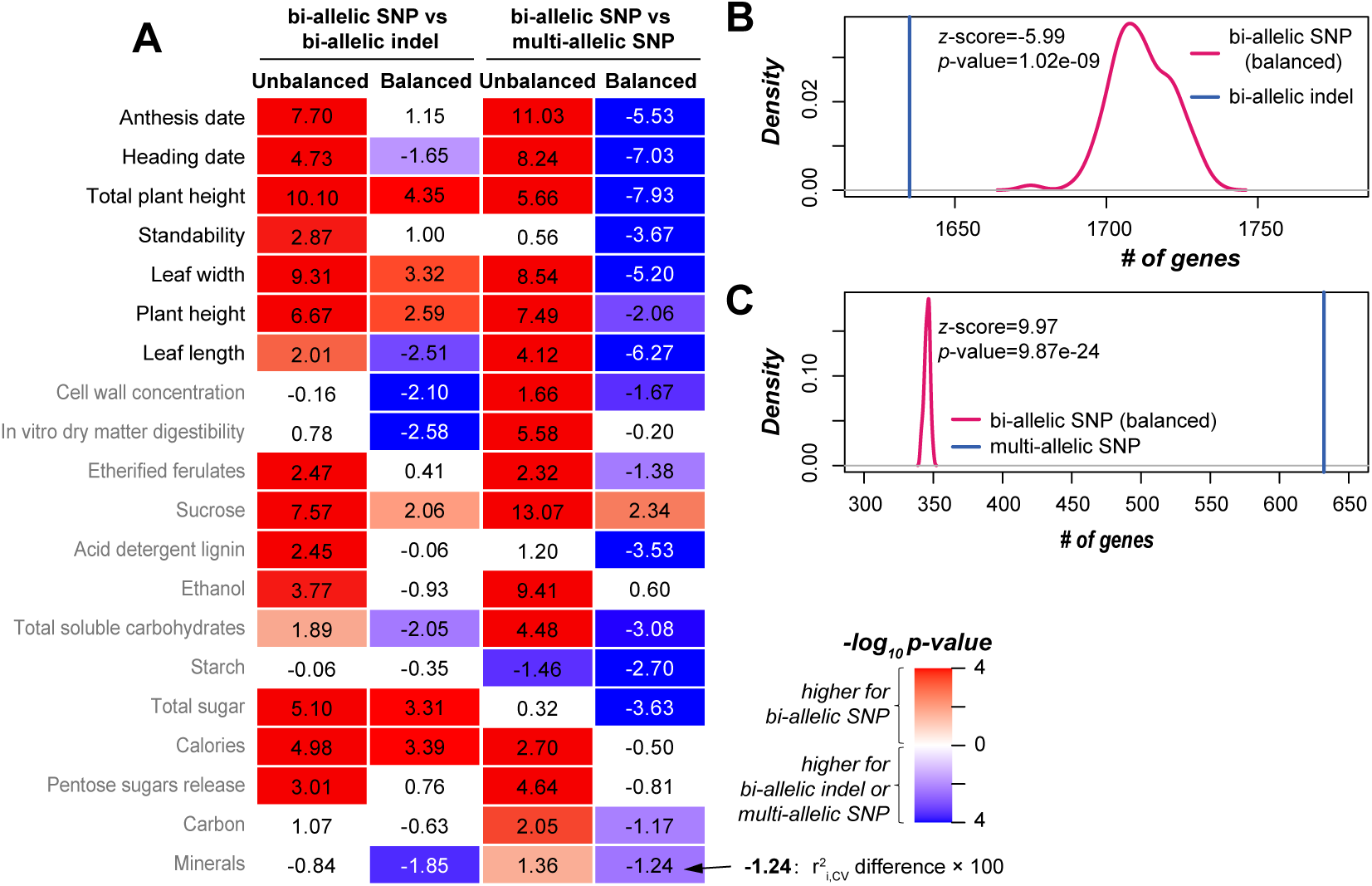
Performance of models built using bi-allelic indels and multi-allelic SNPs. (**A**) Differences in r^2^_i,CV_ between models built using GBS bi-allelic SNPs and models built using GBS bi-allelic indels (left two columns), and between GBS bi-allelic SNP-based models and GBS multi-allelic SNP-based models (right two columns). The “Unbalanced” and “balanced” columns: the bi-allelic SNPs were not and down-sampled to the same numbers of bi-allelic indels or multi- allelic SNPs, respectively. Values in the heatmap: r^2^_i,CV_ differences × 100; only the comparisons with statistical significance (*p* from Wilcoxon signed-rank test < 0.05) are indicated with colors; color scale: -log_10_ *p*-value; red: significantly higher r^2^_i,CV_ was observed for the bi-allelic SNP-based models; blue: significantly higher r^2^_i,CV_ was observed for models built using bi-allelic indels or multi-allelic SNPs. (**B**) Comparison between the numbers of genes harboring or adjacent to the down-sampled bi-allelic SNPs (pink line) and bi-allelic indels (blue line). (**C**) Comparison between the numbers of genes harboring or adjacent to the down-sampled bi-allelic SNPs (pink line) and multi-allelic SNPs (blue line).

In addition to bi-allelic indels, there were 1,836 multi-allelic variants identified for the GBS data, including multi-allelic SNPs (350), multi-allelic indels (604) and multi-allelic SNPs/indels (882) (before encoding, **Table S3**). To simplify the comparison, we built models using only the 350 GBS multi-allelic SNPs. Multi-allelic SNP-based models had significantly lower r^2^_i,CV_ than models based on bi-allelic SNPs (without down-sampling) for 16 traits (the 3^rd^ column in **Fig. 4A)**, but had significantly higher r^2^_i,CV_ for 15 traits than models built using down-sampled bi-allelic SNPs (the 4^th^ column in **Fig. 4A**). Consistent with our earlier results indicating the importance of gene coverage by variants in GP, we found 0.83 times more genes covered by multi-allelic SNPs (632) than balanced bi-allelic SNPs (median: 346, *p*-value = 9.87e-24, **Fig. 4C**), but slightly fewer genes covered by bi-allelic indels (1,635, compared with median 1,711 genes covered by the corresponding balanced bi-allelic SNPs, *p*-value = 1.02e-09, **Fig. 4B**). Taken together, these results indicate that multi-allelic SNPs outperform the commonly used bi-allelic SNPs in trait prediction, particularly in their contribution to improve the coverage of genes, and should be included in future studies.

Next, we asked whether the prediction can be improved by taking advantage of all the types of variants. We built new GP models using all the GBS variants, including 13,265 bi- and multi- allelic SNPs and indels in total. This was not conducted for EC variants, due to the large number of EC variants (2,552,214 variants in total) and the extremely high computing resource requirement to establish models using all the EC variants. We found that five and seven traits had significantly improved r^2^_i,CV_ and r^2^_i,_test in the integrated models compared with models that were built using individual types of GBS variants separately, respectively (**Fig. 5A**,**B** for r^2^_i,text_, r^2^_p_ and r^2^_g_, see **Fig. S8**). These results suggest that different types of genetic variants should be integrated to improve the GP accuracy for switchgrass traits, and potentially traits in other plant species as well.

**Fig. 5.**
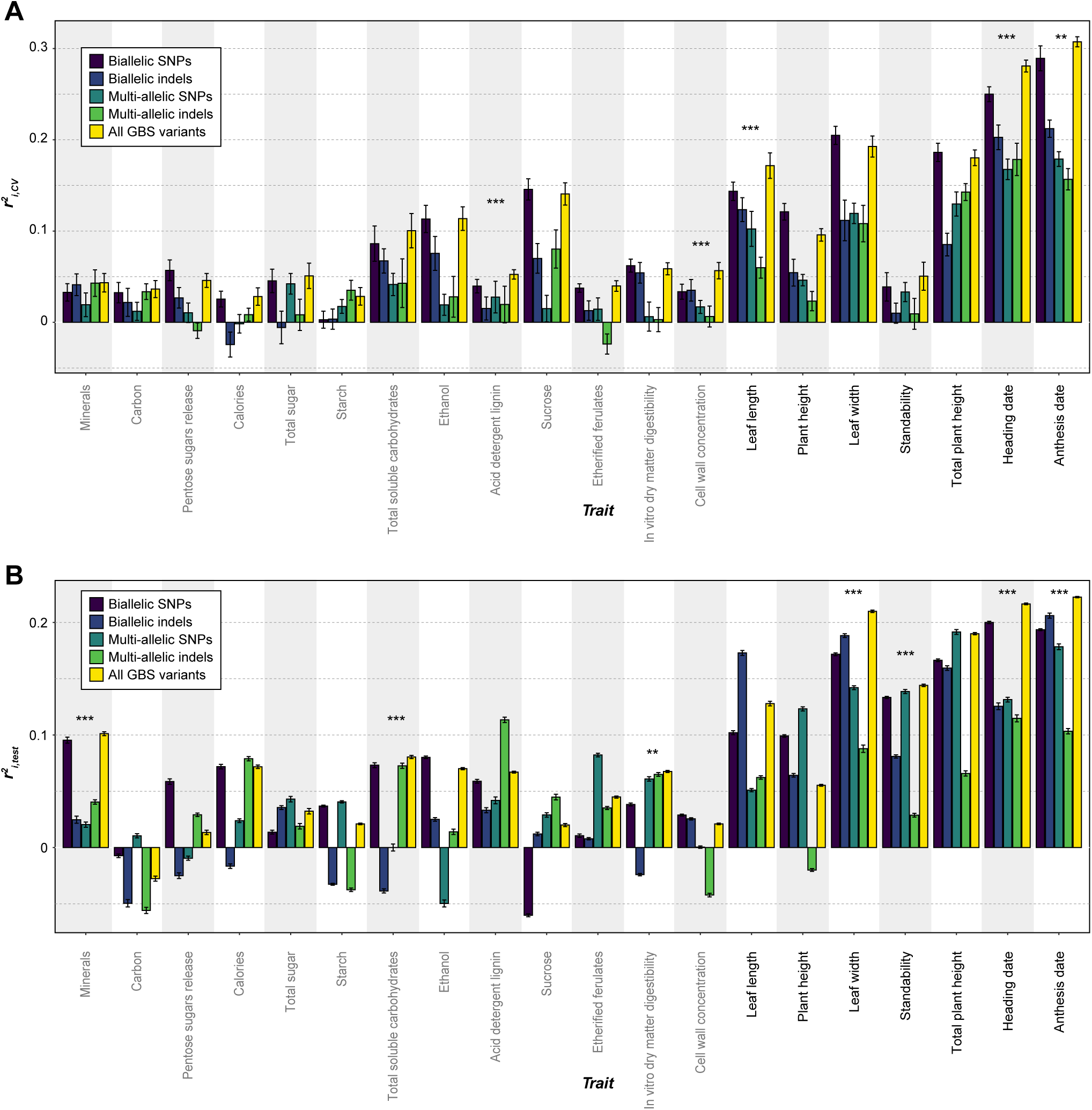
Improvement in prediction accuracy by integrating different types of GBS variants. (**A,B**) The r^2^_i,CV_ (**A**) and r^2^_i,test_ (**B**) for models built using GBS bi-allelic SNPs (dark blue), bi-allelic indels (blue), multi-allelic SNPs (dark green), multi-allelic indels (green), and all the GBS variants (yellow). Column with ** or ***: the integrating model had significantly improved prediction accuracy compared with all the models built using a single type of GBS variant; **: *p*-value from Wilcoxon signed-rank test < 0.01; ***: *p*-value < 0.001. Error bar: standard deviation.

### Models built for octoploids had lower trait prediction accuracy than those for tetraploids

Thus far, the GP models were built for all individuals, including both tetraploids and octoploids, where the variants for octoploid individuals were treated using a pseudo-diploid genotyping strategy (see **Methods**), which is commonly used to handle variants for individuals with different ploidy levels in previous studies. Considering the relatively higher complexity of variants for octoploid individuals, we asked whether the same variant set (GBS or EC) would have different prediction accuracy for tetraploid and octoploid individuals. When GBS bi-allelic SNPs were used, models built for tetraploids had significantly higher r^2^_i,CV_ than octoploid models for 16 traits, but significantly lower r^2^_i,CV_ for only 3 traits (the “GBS” column, **Fig. 6A**). One possible explanation of the lower r^2^_i,CV_ for octoploid models is that the variants called for octoploids were not as good as those for tetraploids in GBS data, due to the relatively low read depth (RD) of GBS bi-allelic SNPs (average at 4.41, left panel in **Fig. 6B**), and potentially a need for deeper RD for octoploids than tetraploids in variant calling. In contrast, the EC data had an average RD at 21.10 (left panel in **Fig. 6C**), and much lower proportions of missing data for both tetraploids (3.1% [16,237], middle panel in **Fig. 6C**, compared with 12.5% [916] for GBS, **Fig. 6B**) and octoploids (2.4% [13,023], right panel in **Fig. 6C**, compared with 12.1% [893] for GBS, **Fig. 6B**). EC-based models for tetraploids still had significantly higher r^2^_i,CV_ than octoploid models for 12 traits (for the GBS-based models, this number was 16), but significantly lower r^2^_i,CV_ for the other 6 traits (3 for the GBS-based models) (the “Exome capture” column, **Fig. 6A**). These results indicate that the low RD in GBS data might, but only partially, explain the lower GP accuracy of octoploid models.

**Fig. 6.**
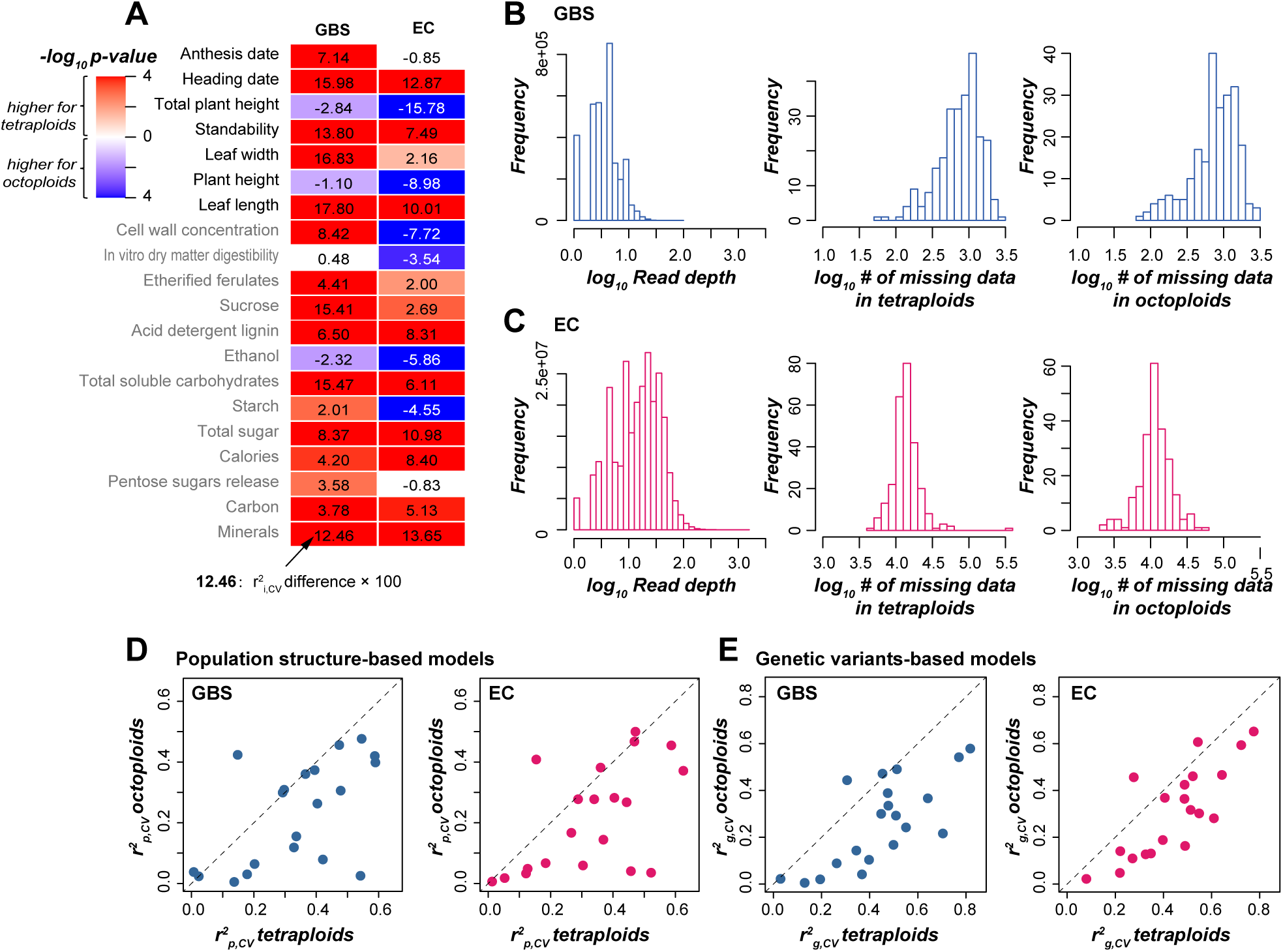
Models for octoploids had poorer prediction accuracy than models for tetraploids. (**A**) Differences in prediction accuracy between models for tetra- and octoploids when GBS and EC bi-allelic SNPs were used. Values in the heatmap: r^2^_i,CV_ differences × 100; only the comparisons with statistical significance (*p* from Wilcoxon signed-rank test < 0.05) are indicated with colors; color scale: -log_10_ *p*-value; red and blue: significantly higher r^2^_i,CV_ was observed for the tetraploid and octoploid model, respectively. (**B**) Read depth (left panel) and the number of missing data (middle and right panels) of the GBS data. (**C**) Read depth (left panel) and the number of missing data (middle and right panels) of the EC data. (**D,E**) Comparison of the r^2^_p,CV_ (**D**) and r^2^_g,CV_ (**E**) between models for tetraploids (x-axis) and octoploids (y-axis) when GBS (left panel) or EC (right panel) bi-allelic SNPs were used.

Another potential explanation for lower GP accuracy for octoploids is the different population structures between octoploids and tetraploids, since both r^2^_p,CV_ (**Fig. 6D**) and r^2^_g,CV_ (**Fig. 6E**) for octoploids were generally smaller than those values for tetraploids, regardless of whether GBS or EC data were used. One previous study (Evans et al., 2015) showed that the within- subpopulation genetic distances in two octoploid subpopulations were shorter than those in three tetraploid subpopulations (see **Fig. 4** in Evans et al., 2015). We further built a model for each of four subpopulations with ≥85 individuals each (**Table S1**) using EC bi-allelic SNPs, and found that subpopulations with larger within-subpopulation genetic distances tended to have higher GP accuracy (either r^2^_p,CV_, r^2^_g,CV_ or r^2^_i,CV_) than those with shorter distances (**Fig. S9**). This is consistent with findings in diverse panels of rice and maize that within-subpopulation genetic variance dominated predictions (Guo et al., 2014), but inconsistent with findings in Barley that GP accuracy decreased with the increase of genetic distances in the training population (Lorenz and Smith, 2015). Taken together, these results indicate that read depth for genetic variants and population structure are potentially another two factors influencing GP accuracy, especially for octoploids in switchgrass.

### Insights of molecular mechanisms underlying trait determination by interpreting GP models

Besides the prediction of complex traits, the GP models can be used to get insights of molecular mechanisms underlying trait variations as well, by examining the absolute coefficients of genetic variants in the trained models. Variants with absolute coefficients ranked above 95^th^ or 99^th^ percentiles were considered as important to trait prediction, and genes harboring or nearby the important variants (<3.5 Kb) were considered as important genes. We focused on the anthesis date models since the current knowledge for genetic bases of flowering time is the most abundant among all the 20 traits studied here, and anthesis date had the highest r^2^_i,CV_ (**Fig. S3**) and one of the highest r^2^_i,test_ (**Fig. S6**). To test how well anthesis date models recover flowering time genes, we collected 23 maize and 378 Arabidopsis flowering time genes (**Table S5**) to identify switchgrass orthologs that were used as benchmark flowering time genes (hereafter referred to as FT-genes, see **Methods**, **Table S6**).

We found that the number of FT-genes identified based on model coefficients is not correlated with model performance (r^2^_i,CV_), either at the 95^th^ (PCC=0.56, *p*-value=0.15) or 99^th^percentile (PCC=0.45, *p*-value=0.27) threshold, suggesting that models with better anthesis date prediction do not necessarily had more FT-genes identified as important. In contrast, the number of FT-genes identified by a model was positively correlated with the number of variants used in the model (**Fig. 7A**, **Fig. S10**, **Table S7-S15**). Specifically, GBS bi-allelic SNPs, bi-allelic indels, multi-allelic SNPs and multi-allelic indels identified only 4 (1), 2 (0), 0 (0) and 1 (0) FT-genes as important, respectively, whereas EC bi-allelic SNPs, bi-allelic indels, multi-allelic SNPs and multi- allelic indels identified 187 (51), 102 (13), 162 (37) and 314 (106) FT-genes as important, respectively, when 95^th^ (99^th^) percentile was used as a threshold (**Fig. 7B**, **Table S15**). Only models built using EC bi-allelic SNPs and EC multi-allelic indels identified significantly more FT- genes than random guessing (*p*-value for fisher’s exact test < 0.034, red bars in **Fig. 7B**) and had the highest odds ratios among other models.

**Fig. 7.**
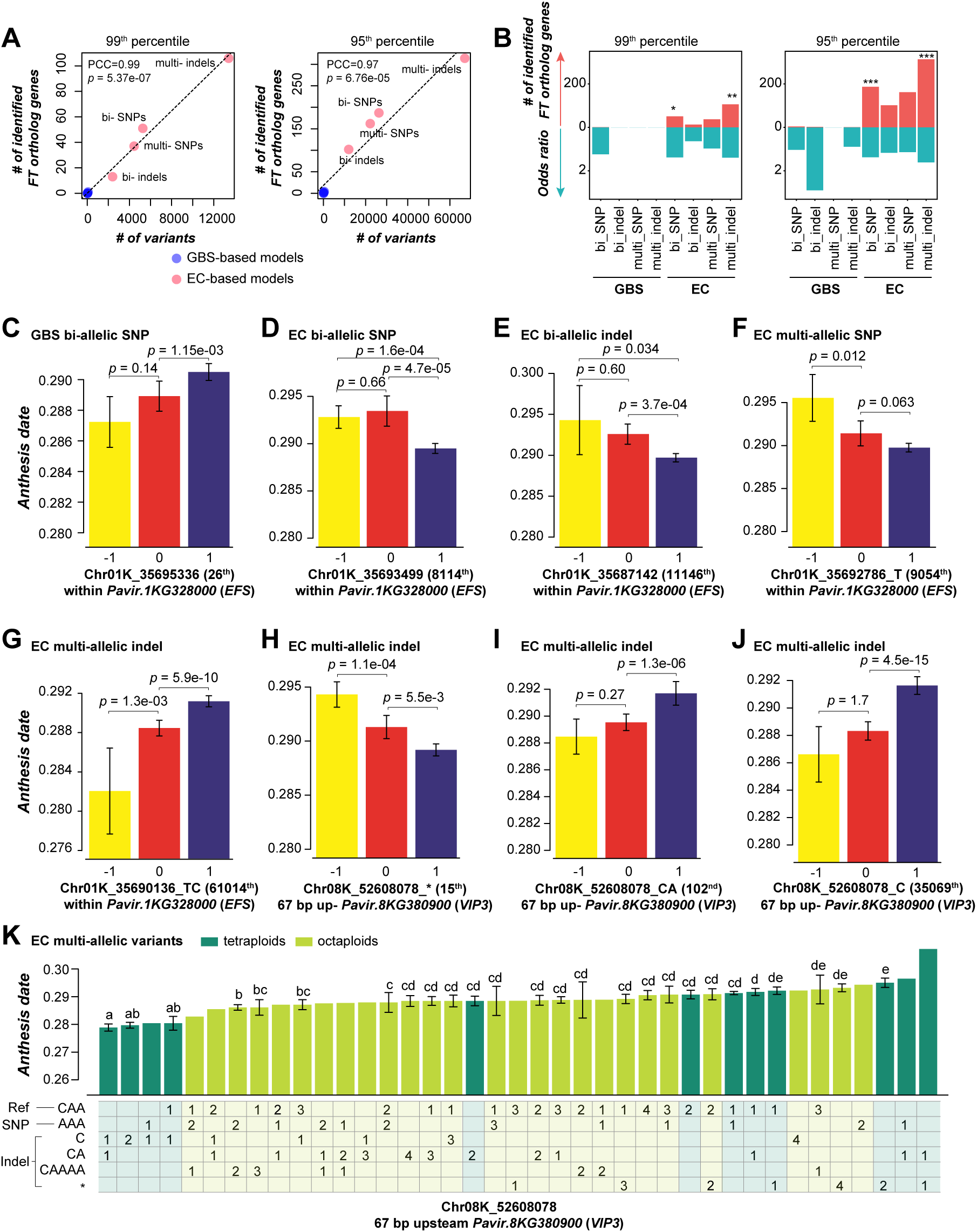
Analysis of important variants for genomic prediction models. (**A**) Correlation between the numbers of variants with absolute coefficient above 99^th^ (left panel) or 95^th^ (right panel) percentile (x-axis) and the corresponding numbers of switchgrass genes orthologous to known flowering time genes (y-axis). Blue: GBS-based models; pink: EC-based models. Dashed line: the diagonal line. (**B**) The number of identified flowering time ortholog genes by models (red bars) and the corresponding odds ratio (green bars) when 99^th^ (left panel) and 95^th^ (right panel) percentile was used as cut-off thresholds. *: *p*-value from Fisher’s exact test < 0.05; **: *p*-value < 0.01; ***: *p*-value < 0.001. (**C-J**) Anthesis dates of individuals with different alleles. The type of variants which were used to build the models were listed above the figures. The absolute coefficient rank of the variant was listed in the parenthesis on the right of the variant. *EFS*: *EARLY FLOWERING IN SHORT DAYS*; *VIP3*: *VERNALIZATION INDEPENDENCE 3*. Yellow, red, and blue bars in (**C-J**): individuals with homologous alternative (-1), heterozygous (0) and homologous reference (1) alleles, respectively. *p*-values were from Wilcoxon signed-rank test. In (**K**), six types of alleles for the locus Chr08K_52608078 were indicated on the left of the table; values in the table indicate the numbers of the corresponding alleles; green: tetraploids; prasinous: octoploids; error bar: standard error; column with no error bar: there was only one individual containing the corresponding allele composition. Only columns with > 1 individual are indicated with letters indicating the statistical significance.

Out of the seven FT-genes identified by GBS variants as important genes when the 95^th^ percentile was used as the threshold, six were also identified by EC variants (**Fig. S10, Table S15**). For example, the GBS bi-allelic SNP Chr01K_35695336, which ranked 26^th^ in terms of absolute coefficient, was located within the genic region of *Pavir.1KG328000*, an ortholog of Arabidopsis *EARLY FLOWERING IN SHORT DAYS* (*EFS*), which represses flowering in the autonomous promotion pathway (Soppe *et al*., 1999). Within *Pavir.1KG328000*, there were also several EC bi-allelic SNPs (e.g., Chr01K_35693499), bi-allelic indels (Chr01K_35687142), multi- allelic SNPs (Chr01K_35692786_T) and multi-allelic indels (Chr01K_35690136_TC) with absolute coefficients ranking above 95^th^ percentile in the corresponding models. Individuals with homozygous reference alleles (1) for these loci flowered significantly differently as individuals with homozygous alternative (-1) and heterozygous alleles (0) (**Fig. 7C-G**). Besides the seven FT-genes identified by GBS variants, all the other identified FT-genes were EC variant-specific, further suggesting that EC variants are superior to GBS variants in either trait prediction or identification of potential contributing genes for traits in question.

Furthermore, we investigated in detail how multi-allelic variants contributed to trait prediction, by taking the locus Chr08K_52608078 as an example. This locus was located 67 bp upstream of *Pavir.8KG380900*, an ortholog of the Arabidopsis flowering time gene *VERNALIZATION INDEPENDENCE 3* (*VIP3*), which functions as an activator of the flowering-repressor gene *FLOWERING LOCUS C* (*FLC*) (Zhang et al., 2003). This locus had six alleles in the EC data among individuals studied here: the reference allele is CAA, and five alternative alleles included one SNP: AAA, and four indels: C, CA, CAAAA, and * (CAA was completely absent). In the model built using EC multi-allelic indels, Chr08K_52608078_* (ranked 15^th^, **Fig. 7H**), Chr08K_52608078_CA (ranked 102^nd^, **Fig. 7I**), and Chr08K_52608078_C (ranked 35069^th^, **Fig. 7J**) had absolute coefficients above the 95^th^ percentile, and individuals with different alleles of these three indels had significantly different anthesis date. Interestingly, when examining different alternative alleles for this single locus, individuals with homozygous reference alleles (see **Methods**) flowered significantly earlier (**Fig. 7H**) or later (**Fig. 7I,J**) than individuals with heterozygous alleles and homozygous alternative alleles, suggestive of the complexity of multi- allelic variants in trait determination. In addition, when examining the original allele compositions of this locus among individuals (**Fig. 7K**), we found that there were 39 combinations of all the six alleles. Tetraploid individuals with homozygous or heterozygous alleles of C flowered earliest, and tetraploid individuals with */*, AAA/CA and CA/* flowered latest, whereas all the other tetraploids and all the octoploids flowered medially. This finding suggests that promoter sequences with a C at this locus may interrupt the expression of *Pavir.8KG380900*, leading to decreased expression of orthologs of *FLC* and resulting earlier flowering phenotypes, if the regulatory rules of *VIP3* and *FLC*-like genes in flowering are conserved between Arabidopsis and switchgrass. This type of analysis highlights the valuable insights into trait prediction provided by multi-allelic variants that were normally neglected by previous studies, and the need to include multi-allelic variants in future studies of GWAS and GP.

## Discussion

Using switchgrass as a model system, our study showed the differences of two types of variants (e.g., GBS vs EC SNPs) for a given factor (e.g., genotyping approach) in the prediction accuracy for 20 traits, and proposed suggestions that can be potentially taken in future GP practices. Generally, different versions of genome assemblies (short read-based vs. long read- based) had similar GP accuracy; genotyping approaches that capture variants with higher gene coverages should be considered with limited budgets; multi-allelic variants should be included in GP practices both for improving GP accuracy and identifying potential contributing variants for the target traits; and different types of variants should be integrated to improve the GP accuracy.

Although these conclusions summarized above were generally true for most traits, there were a few exceptions. For example, EC-based models tended to outperform GBS-based models for most traits, but the plant height (base of longest flowering stem to the node at the base of the panicle) and total plant height (base of the longest flowering stem to the tip of the panicle) were predicted with consistently higher accuracy by GBS-based models, no matter whether the balanced, unbalanced, genic or intergenic variants were used (**Fig. 3A**). This finding indicates that variations in plant height might be determined more by variants in genomic elements that are located within intergenic regions than those within genic regions. In addition, models built for tetraploids tended to have better prediction accuracy for most traits than those for octoploids, but the two plant height traits and ethanal (ethanol/g dry forage) were better predicted in octoploid models, both when GBS and EC variants were used (**Fig. 5A**). These exceptions, together with findings in Wang et al. (2023a) that whether Deep Learning approaches outperformed classical approaches depends on the datasets used and traits in question, suggest that the optimal GP practices should be considered with caution on a case-by-case basis.

Based on our findings, in switchgrass, short read-based assembly led to comparable (for 12 traits) or even significantly better prediction accuracy (for 7 traits) for switchgrass complex traits, whereas long read-based assembly only led to better prediction accuracy for anthesis date, for which the difference in r^2^_i,CV_ between short read- and long read-based models was only 0.0163. Those results suggest that, for a plant species or lineage with only reference genome assembly based on short reads, it might not be necessary to update the assembly using long reads if only for improving the GP accuracy. However, if both the short and long read-based assemblies are available, long read-based assembly will still be a better option since ⅰ) the differences in r^2^_i,CV_ were relatively small (max = 0.0324); ⅱ) long read-based assembly tend to be more commonly used by the community, and is much more competent than short read-based assembly for calling SVs, another genetic variants that were not studied in this study but have been shown great power in GP and GWAS practices.

In addition, our findings suggested that multi-allelic variants outperformed bi-allelic ones in trait prediction and identification of potential important genes that contribute to the formation of traits in question. Multi-allelic tandem repeats (He et al., 2024) and multi-allelic copy number variations (Handsaker et al., 2015) have been shown to contribute to the rice agronomic traits and gene expression dosage variations in humans, respectively. Although ∼10% (Jiang et al., 2020) or even a higher proportion (Kang et al., 2023) of genetic variants are multi-allelic, previous studies generally used bi-allelic variants only. Besides the strategy we used in this study to encode the multi-allelic variants, the BCFtools offered a model recently to handle loci with multiple alternative alleles (Danecek et al., 2021), which would help us make the best use of all the genetic variants in GWAS and GP practices.

Our results indicate that gene coverage by variants had a significant effect on GP accuracy, which gives recommendations for the selection of genotyping approaches and designation of SNP chips in future studies. A pan-genome study of 69 *A. thaliana* accessions showed that 18% of the 32,986 gene families were private to a single accession (Lian et al., 2024), indicating potential missing heritability by using a single reference genome. However, in most previous studies and our study, genetic variants were generally called by referencing a single genome assembly. Recently, more and more graph-based pan-genomes have been constructed to capture the missing heritability in plant species (Zhou et al., 2022; Kang et al., 2023; Yan et al., 2023; Liu et al., 2024). For example, including SVs identified based on pan-genome has been shown to significantly improve the estimated heritability for grape traits (Liu et al., 2024). It is important to know whether including other hidden genomic variants (e.g., SNPs and indels) revealed by pan-genome would also help improve GP accuracy for plant traits, especially for traits enabling a small group of ecotypes/cultivars/individuals to adapt to particular environments or endowing them certain horticultural characteristics to be distinguished from others.

In this study we showed that models built for tetraploids tended to have better prediction accuracy for most traits than those for octoploids, potentially due to relatively longer genetic distances among tetraploids. Another potential reason that can not be ruled out is that the allele dosage in octoploids were much more complex than those in tetraploids, whereas we took the strategy of pseudo-diploid genotyping that grouped all heterozygotes (e.g., AT, ATTT, AATT, AAAT) into a single class. Some studies treated heterozygotes in an allele-dosage way (Yadav et al., 2024) or under different effect assumptions (namely additive, duplex dominant, and simplex dominant, Rosyara et al., 2016; Wilson et al., 2021) when dealing with variants for polyploids. Considering that it is unlikely that genes/loci with different allele compositions across the whole genome have a same effect pattern, and that multiple ploidy levels may coexist in some crops (e.g., diploids, triploids, tetraploids and aneuploids coexist in *Phalaenopsis* orchids), tools that can detect the potential effect pattern for each locus based on prior knowledges or integrate all the possible effect patterns of alleles, and handle with multiple levels of ploidies simultaneously, are needed for crop selective breeding. The prior knowledge can be that, under which assumption, the statistical significance between genetic variants and traits of individuals with different allele compositions is the highest.

## Materials and Methods

### Genomic and phenomic data

Two versions of switchgrass genome assemblies (v1 and v5) were downloaded from Phytozome (https://phytozome.jgi.doe.gov/pz/portal.html). The GBS data was obtained from (Lu et al., 2013), and the raw data were downloaded from NCBI under the project PRJNA201059 (**Table S1**). The GBS barcodes for individuals in each pooled GBS library were obtained from the description of each SRA sample in the NCBI website. The EC data was from (Evans et al., 2015), and the raw data were downloaded from NCBI under the project PRJNA280418 (**Table S1**). Phenotypic data were from the diversity panel which consisted of 540 individuals from 66 populations (Lipka et al., 2014), including seven morphological and 13 biochemical traits (**Table S2**), where the cumulative growing degree days was used to record the anthesis date and heading date. After filtering out individuals with low quality of GBS data, ploidy of 6, and the ones with information missing from any of the 20 traits, we retained 486 individuals in this study (**Table S2**), including 263 tetraploids and 223 octoploids.

### SNP calling

Reads from the GBS and EC data were trimmed using Trimmomatic (Bolger et al., 2014) with parameters: 2:30:10 LEADING:3 TRAILING:3 SLIDINGWINDOW:4:20 MINLEN:35. Trimmed reads with scores > 20 and longer than 35 bp were kept for the following analysis. GBS reads were mapped to both two assemblies using bwa/0.7.12.r1044 (Li, 2013) with default parameter setting, while EC reads were only mapped to the v5 assembly. The alignment after mapping was sorted using picardTools/1.113 (http://broadinstitute.github.io/picard).

SNP calling was conducted using GATK/3.5.0 (Auwera and O’Connor, 2020). For EC data, duplication alignments, which may be amplified during library preparation progress, were marked using MarkDuplicates module. No treatment was processed to GBS alignments, because reads in GBS data always had the same start point (the same restriction size, *Ape*KI site) (Lu et al., 2013) and stop point. SNPs and indels were identified using HaplotypeCaller module, with parameters of -stand_call_conf 40 -stand_emit_conf 10 --max_alternate_alleles 8 -ploidy x, where x = 2 for tetraploids (since the reference genome was also a tetraploid) while 4 for octoploids. Base-quality recalibration was conducted using BaseRecalibrator and IndelRealigner for SNPs and indels, respectively. Then the HaplotypeCaller module was used for the second run of SNP and indel calling with the same parameter setting and the output files were saved at GVCF format. All GVCF files for individuals were merged together using the CombineGVCFs module. The resulting SNPs were filtered using VariantFiltration with parameters of “QD < 2.0 || MQ < 40.0 || FS > 60.0 || MQRankSum < –12.5 || ReadPosRankSum < –8.0 –clusterSize 3 – clusterWindowSize 10”, while the indels were filtered with “QD < 2.0 || FS > 200.0 || ReadPosRankSum < –20.0”. Finally, SNPs and indels with minor allele frequency (MAF) > 0.05, and missing data < 20% were kept in the final list.

### Encoding of multi-allelic variants

Multi-allelic variants were encoded using the method proposed by Zhan et al. (2016)with modifications, where m columns were encoded for m alternative alleles (m≥1). For example, for a tetra-allelic variant (e.g, the reference allele is A, three alternative alleles are T, G, AT), there would be three columns (**Table 1**): A/A, A/T or T/T for the first alternative allele, A/A, A/G or G/G for the second, and A/A, A/AT/ or AT/AT for the third. The first two columns were used as multi- allelic SNPs, and the third column was used as a multi-allelic indel. After encoding the multi- allelic variants, there were 1,628 GBS multi-allelic SNPs, 2,454 GBS multi-allelic indels, 444,102 EC multi-allelic SNPs, and 1,341,337 EC multi-allelic indels.

**Table 1.** Encoding for multi-allelic variants with A as the reference allele, and T, G and AT as the alternative alleles.

|  | Sequence | Column_1 (T) | Column_2 (G) | Column_3 (AT) |
| --- | --- | --- | --- | --- |
| ID_1 | A/A | 1 (A/A) | 1 (A/A) | 1 (A/A) |
| ID_2 | A/T | 0 (A/T) | 1 (A/A) | 1 (A/A) |
| ID_3 | T/G | 0 (A/T) | 0 (A/G) | 1 (A/A) |
| ID_4 | G/AT | 1 (A/A) | 0 (A/G) | 1 (A/AT) |
| ID_5 | AT/AT | 1 (A/A) | 1 (A/A) | -1 (AT/AT) |

### Imputation of missing data

For bi-allelic SNPs in tetraploids, the missing data were imputed using fastPHASE (Scheet and Stephens, 2006). Variants called for octoploids were treated as those called for tetraploids. For example, a variant with homozygous reference allele AAAA and a variant with homozygous alternative allele TTTT were treated as AA and TT, respectively, while all three types of heterozygous alleles ー ATTT, AATT and AAAT ー were treated as AT. For the bi-allelic indels, the reference allele was converted to “R”, and the alternative allele was converted to “A”, and then the missing data were imputed using fastPHASE. The multi-allelic variants were first encoded to bi-allelic ones, and then were imputed using fastPHASE.

### Genomic prediction model

To build GP models, homozygous reference alleles and alternative alleles were encoded as 1 and -1, respectively, while heterozygous variants were encoded as 0. The R package “rrBLUP” (Endelman, 2011) was used to build the GP models because it is one of the methods which have relatively higher prediction accuracy compared with others in general (Lipka et al., 2014; Azodi et al., 2019). For each trait, all the 486 individuals were first split into training (389, 80%) and test (97, 20%) sets using the stratified strategies to make sure that individuals in training and test sets had similar trait value distribution. The training set was then used to establish predictive models, using a five-fold cross-validation scheme: (1) all the training individuals were first split into five folds; (2) individuals in four folds (referred to as training subset) were used to build models, while individuals in the other fold (referred to as validation subset) was used to evaluate the model performance; (3) the second step was conducted five times to make sure each fold will be used as validation subset once; (4) the r^2^ of PCC between true and predicted trait values of individuals in all five folds was calculated and was referred to as r^2^_CV_. This five-fold cross-validation scheme was repeated 10 times, and the median r^2^_CV_ value of 10 replicate runs was calculated. During the model training, the coefficient of each variant was calculated for each cross-validation step and was averaged across five cross-validations. Then the absolute mean coefficient of a variant was used as the measure of importance for this variant in a model.

To balance the numbers of variants used between two models (e.g., models built using GBS bi-allelic SNPs vs those using EC bi-allelic SNPs), the one with more variants (i.e., EC bi-allelic SNPs) was down sampled to the same number as the one with fewer variants (GBS bi-allelic SNPs). This random down-sampling was conducted 100 times, resulting in 100 down-sampled data. The median prediction accuracy across the 100 models building using these 100 down- sampled data was used to measure the model performance.

### Association of GBS bi-allelic SNPs between two assemblies

To assess which GBS bi-allelic SNPs were shared between v1 and v5 assemblies, two approaches were explored in this study. First, bi-allelic SNPs from two assemblies were associated by aligning sequences of these two assemblies using MUMmer/4.0.0beta2 (Kurtz et al., 2004). Sequences of the v1 chromosomes and contigs were aligned with those of v5 chromosomes and scaffolds, using the function “NUCmer” with the option of --maxmatch. The function “mummerplot” was used to output the coordinates and the sequence identities of matched regions. Only regions with sequence identity > 95% were kept for downstream analysis. For each pair of matched regions, if the v1 region contains a SNP, then search for SNPs on the corresponding v5 region. A v1 SNP was associated with a v5 SNP if sequences 20bp upstream or downstream of the v1 SNP matched to those of the v5 SNP (**Table S4**).

The second approach is to align the v1 SNP in question and its neighbor sequences (500 bp up- and 500 bp downstream) to the v5 genome sequence using BLASTN (Altschul et al., 1990). If the best matched v5 region for a v1 1001 bp region contains a SNP at the same relative coordinate as the v1 SNP, then the v1 SNP and the v5 SNP are associated.

By comparing results from these two approaches, out of 7556 v1 bi-allelic SNPs, 1502 (19.9%) were associated with v5 SNPs from both approaches, 3777 (50.0%) and four (0.05%) were associated with v5 SNPs from only blastn and Mummer methods, respectively. The corresponding v5 sequences of 2118 v1 SNPs (28%) were not identified as having SNPs. This may be because SNPs were filtered by applying a hard threshold on continuous scores (i.e., parameter setting during SNP calling), thus SNPs with scores close to the threshold may be filtered out in one assembly but not in the other one. The flank regions of five v1 SNPs had no matched sequences in v5, 83 v1 SNPs had no matches in v5 assembly near the SNPs, and 67 had different alleles between v1 and v5 assemblies (**Table S4**).

### Orthologs to known flowering time genes

Known genes involved in flowering time in maize were gained from (Azodi et al., 2020) and the Maize Genetics and Genomics Database (https://www.maizegdb.org/). Flowering time genes in *Arabidopsis thaliana* were gained from the Flowering Interactive Database (http://www.phytosystems.ulg.ac.be/florid/, Bouché *et al*. 2016). Orthologs in switchgrass to these known flowering time genes were identified using the software OrthoFinder (Emms and Kelly, 2019) with the default settings. All the known flowering time genes and their orthologs in switchgrass are shown in **Table S5,S6**. Genetic variants that were located in intergenic regions were associated with the nearby genes which are closest to the variants and with distances to the variants < 3.5 Kb.

## Acknowledgments

This work was supported by the U.S. Department of Energy Great Lakes Bioenergy Research Center (BER DE-SC0018409 to SHS), the National Science Foundation (DGE- 1828149 to SHS and KSA; IOS- 2107215, IOS-2218206, and MCB-2210431 to SHS), the National Natural Science Foundation of China (32370241 to PW) and the Scientific Research Foundation of and the Major Scientific Research Tasks from Kunpeng Institute of Modern Agriculture at Foshan (KIMAQD2022003 and KIMA-ZDKY2022004 to PW).

## Author contributions

PW and SHS conceived and designed this study. MDC provided data and advice on study design and interpretation. PW and FM conducted the read mapping and SNP/indel calling analyses. PW conducted all the other analyses with help from CBA and KSA. PW and SHS wrote the manuscript with inputs from all authors. All authors read and approved the final manuscript.

## Data availability

All the scripts used in this study are available on Github at: https://github.com/ShiuLab/Manuscript_Code/tree/master/2022_GP_in_Switchgrass.

---

**Fig. S1.**
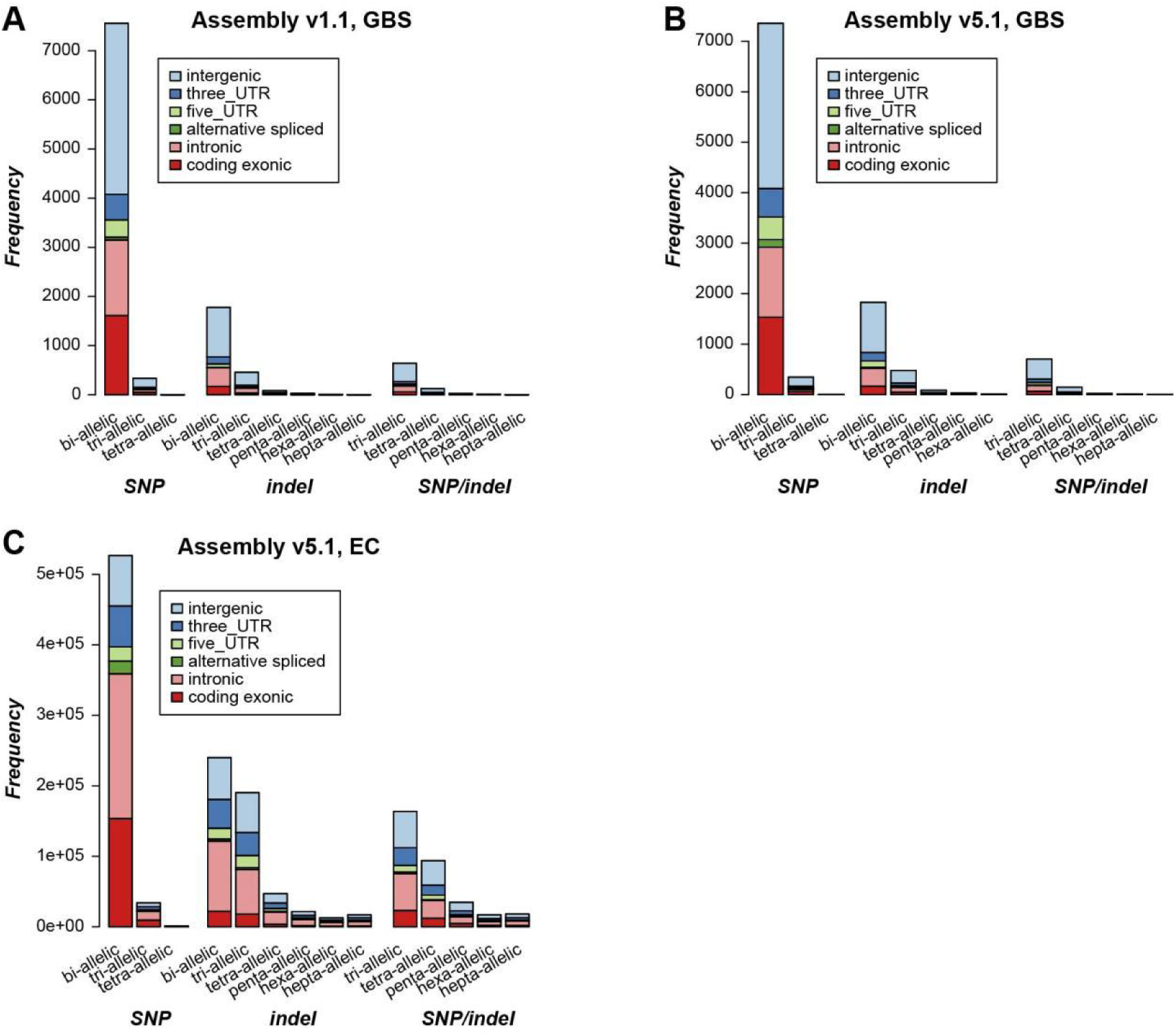
Classification of different types of genetic variants. (**A**) Frequency of different types of variants called by mapping the GBS data to v1 assembly. (**B**) Frequency of different types of variants called by mapping the GBS data to v5 assembly. (**C**) Frequency of different types of variants called by mapping the EC data to v5 assembly.

**Fig. S2.**
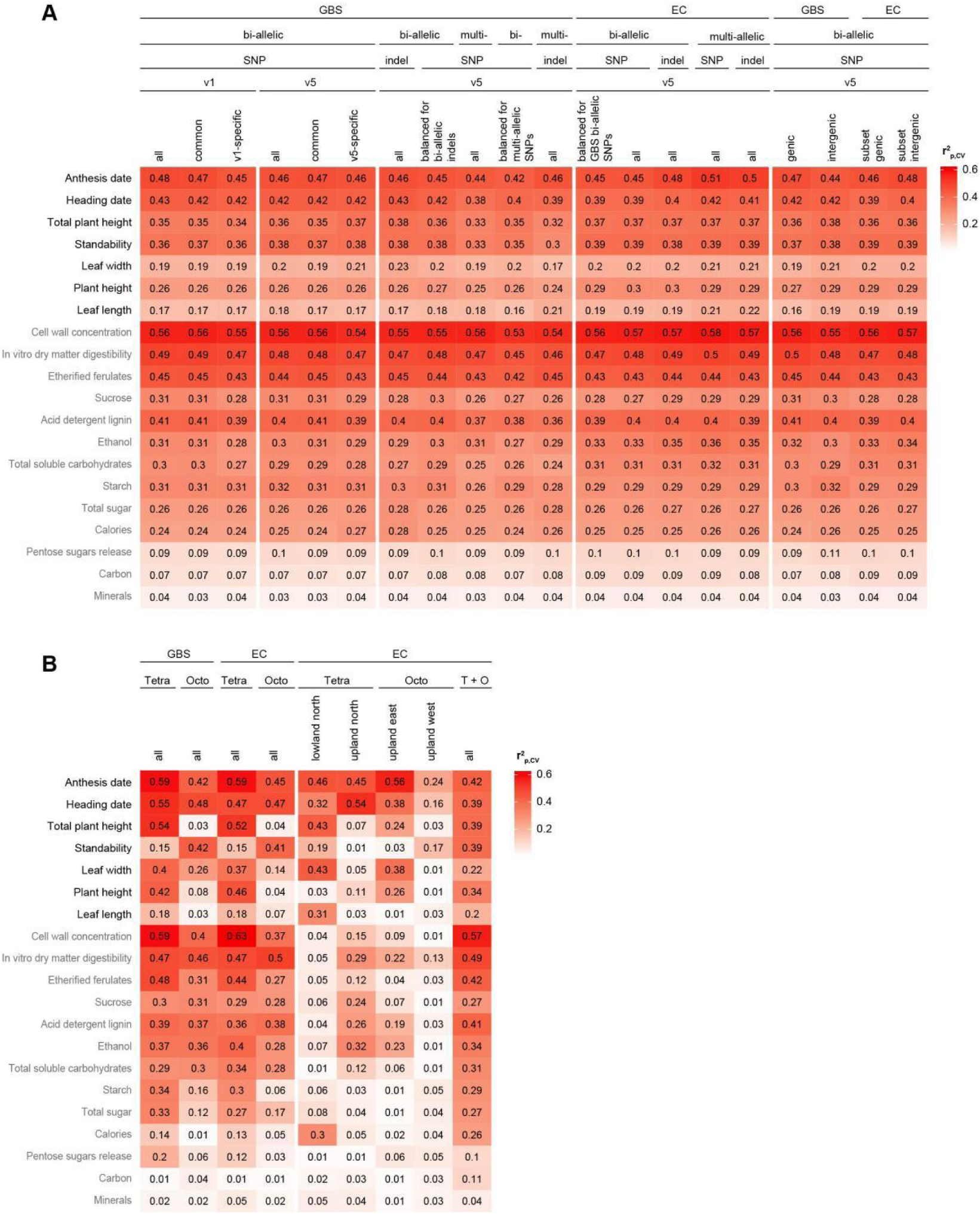
Prediction accuracy of models built using the population structure on the cross- validation sets. (**A**) r^2^_p,CV_ for models built on both tetraploids and octoploids using different types of variants. Table above the heatmap shows different factors the variants used may belong to. For example, the 1^st^ and 4^th^ column show r^2^_p,CV_ of models built using bi-allelic SNPs called by mapping the GBS data to the v1 and v5 assembly, respectively; the 12^th^ and 13^th^ column show r^2^_p,CV_ of models built using balanced subset (down-sampled to the same number of GBS bi-allelic SNPs) and all the v5-based EC bi-allelic SNPs, respectively. (**B**) r^2^_p,CV_ for models built for different ploidy levels or subpopulations using GBS or EC bi-allelic SNPs. Color scale in the heatmap: median r^2^_p,CV_ among 10 replicate runs.

**Fig. S3.**
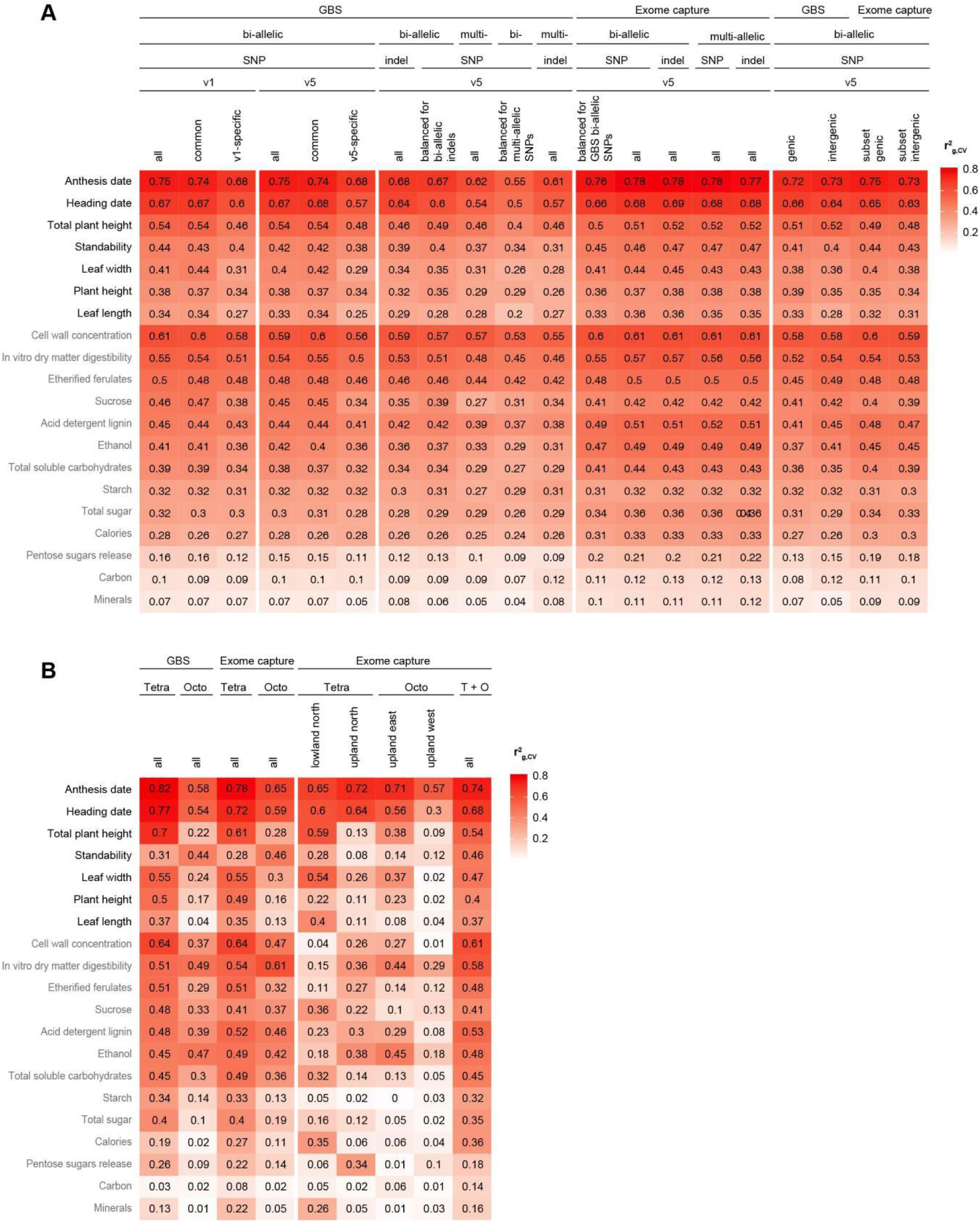
Prediction accuracy of models built using genetic variants on the cross-validation sets. (**A**) r^2^_g,CV_ for models built on both tetraploids and octoploids using different types of variants. Table above the heatmap shows different factors the variants used may belong to. (**B**) r^2^_g,CV_ for models built for different ploidy levels or subpopulations using GBS or EC bi-allelic SNPs. Color scale in the heatmap: median r^2^_g,CV_ among 10 replicate runs.

**Fig. S4.**
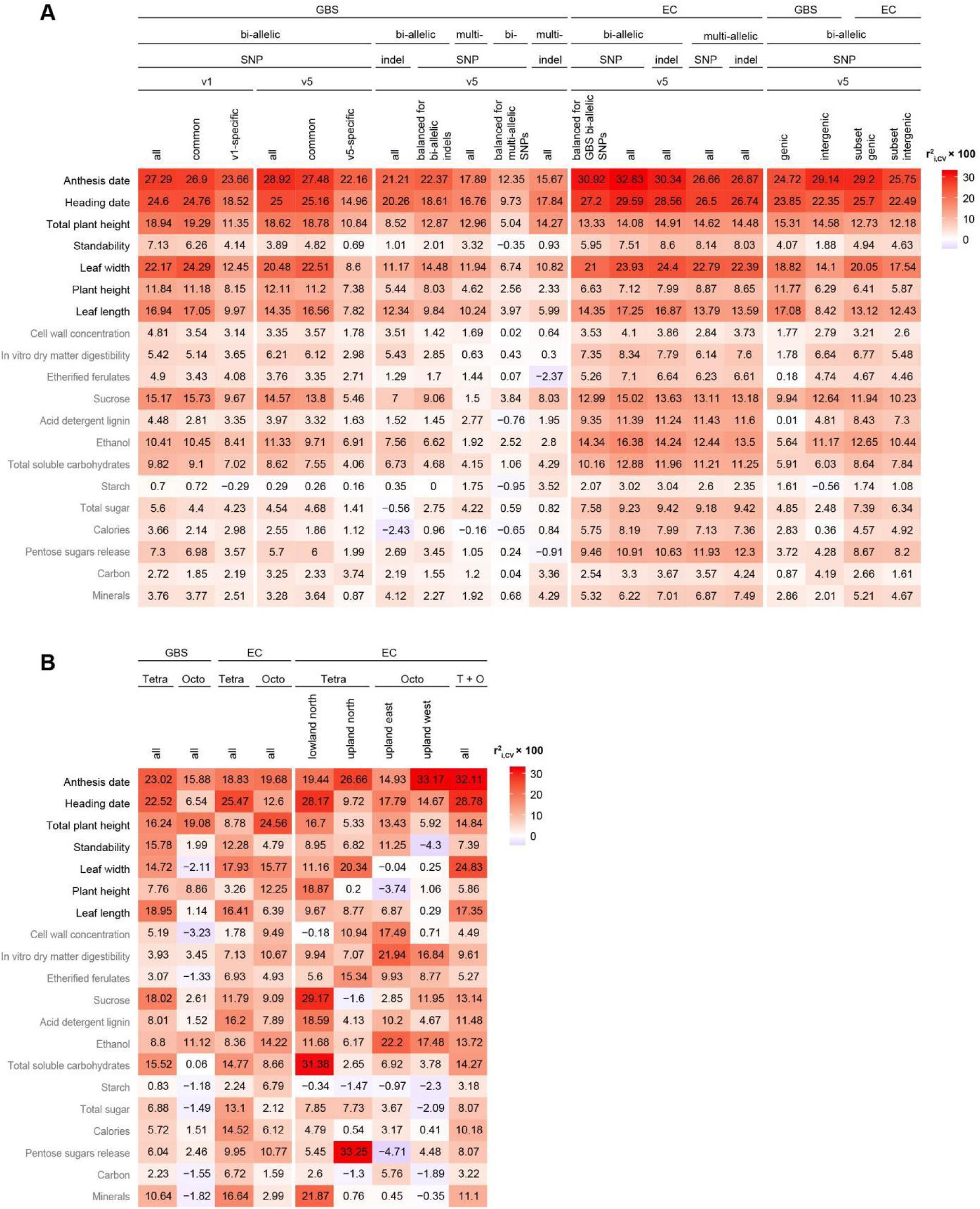
Improvement of r^2^ of models built using different genetic variants on the cross- validation sets. (**A**) r^2^_i,CV_ for models built on both tetraploids and octoploids using different types of variants. Table above the heatmap shows different factors the variants used may belong to. (**B**) r^2^_i,CV_ for models built for different ploidy levels or subpopulations using GBS or EC bi-allelic SNPs. Color scale in the heatmap: r^2^_i,CV_ × 100.

**Fig. S5.**
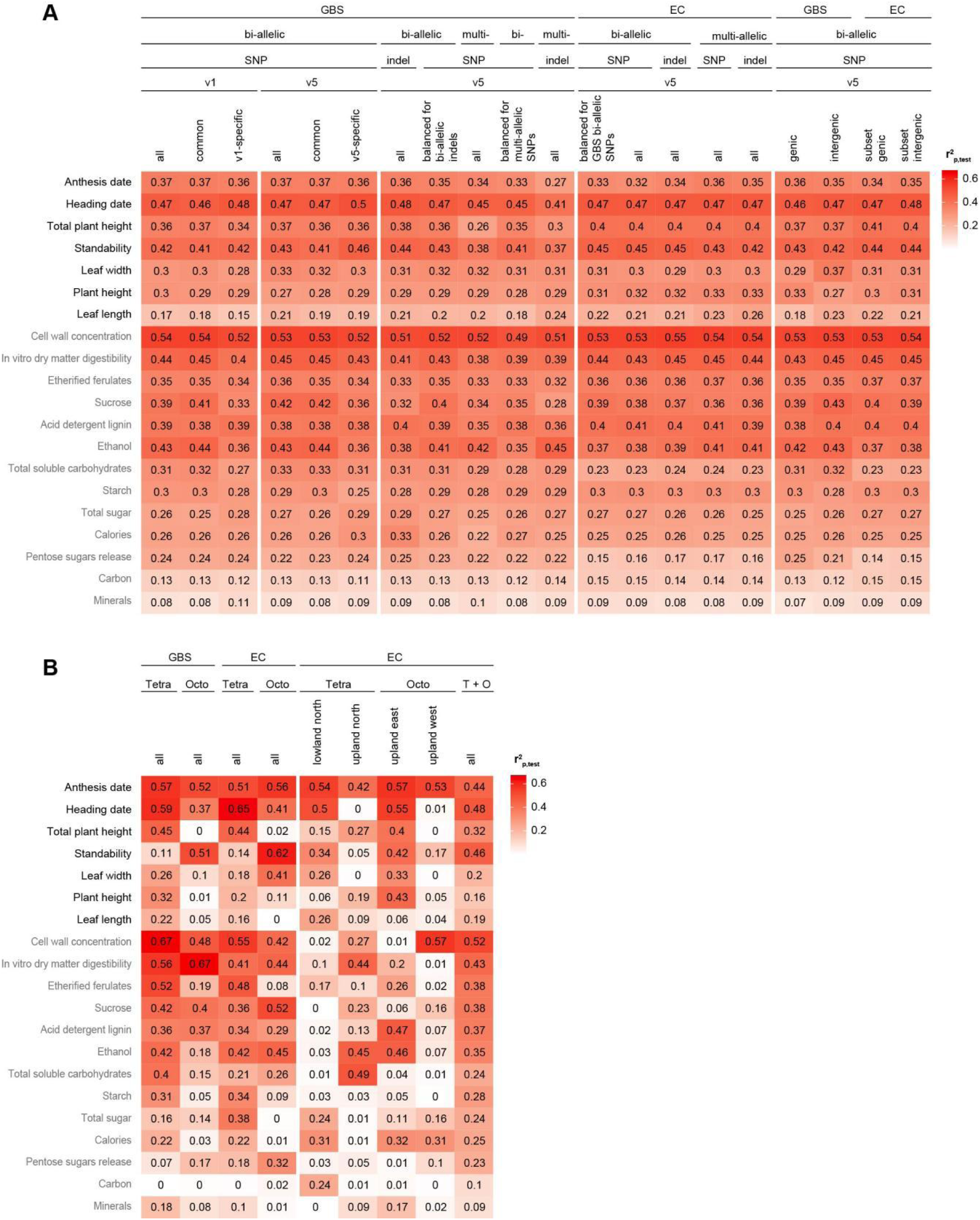
Prediction accuracy of models built using the population structure on the test sets. (**A**) r^2^_p,test_ for models built on both tetraploids and octoploids using different types of variants. Table above the heatmap shows different factors the variants used may belong to. (**B**) r^2^_p,test_ for models built for different ploidy levels or subpopulations using GBS or EC bi-allelic SNPs. Color scale in the heatmap: median r^2^_p,test_ among 10 replicate runs.

**Fig. S6.**
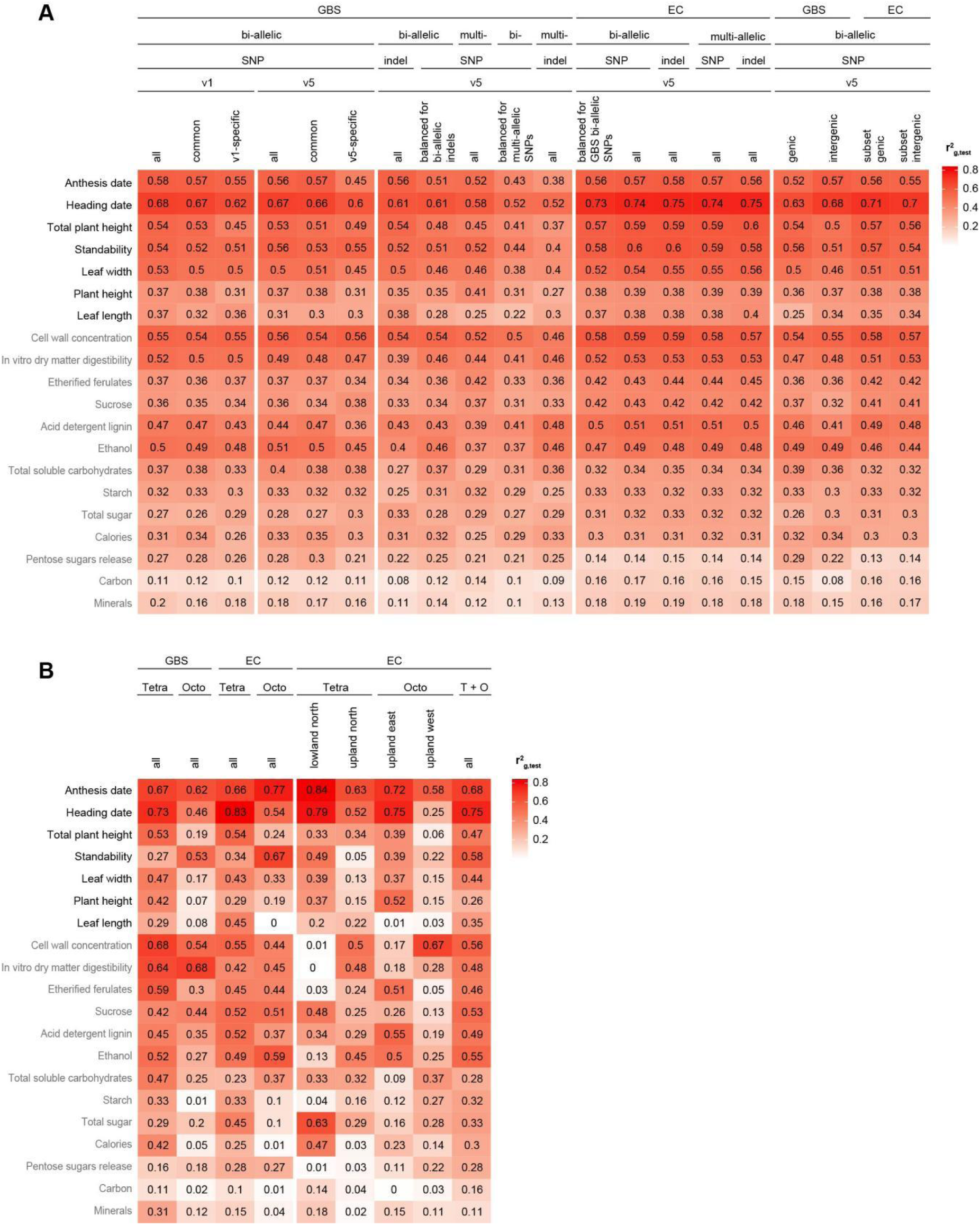
Prediction accuracy of models built using genetic variants on the test sets. (**A**) r^2^_g,test_ for models built on both tetraploids and octoploids using different types of variants. Table above the heatmap shows different factors the variants used may belong to. (**B**) r^2^_g,test_ for models built for different ploidy levels or subpopulations using GBS or EC bi-allelic SNPs. Color scale in the heatmap: median r^2^_g,test_ among 10 replicate runs.

**Fig. S7.**
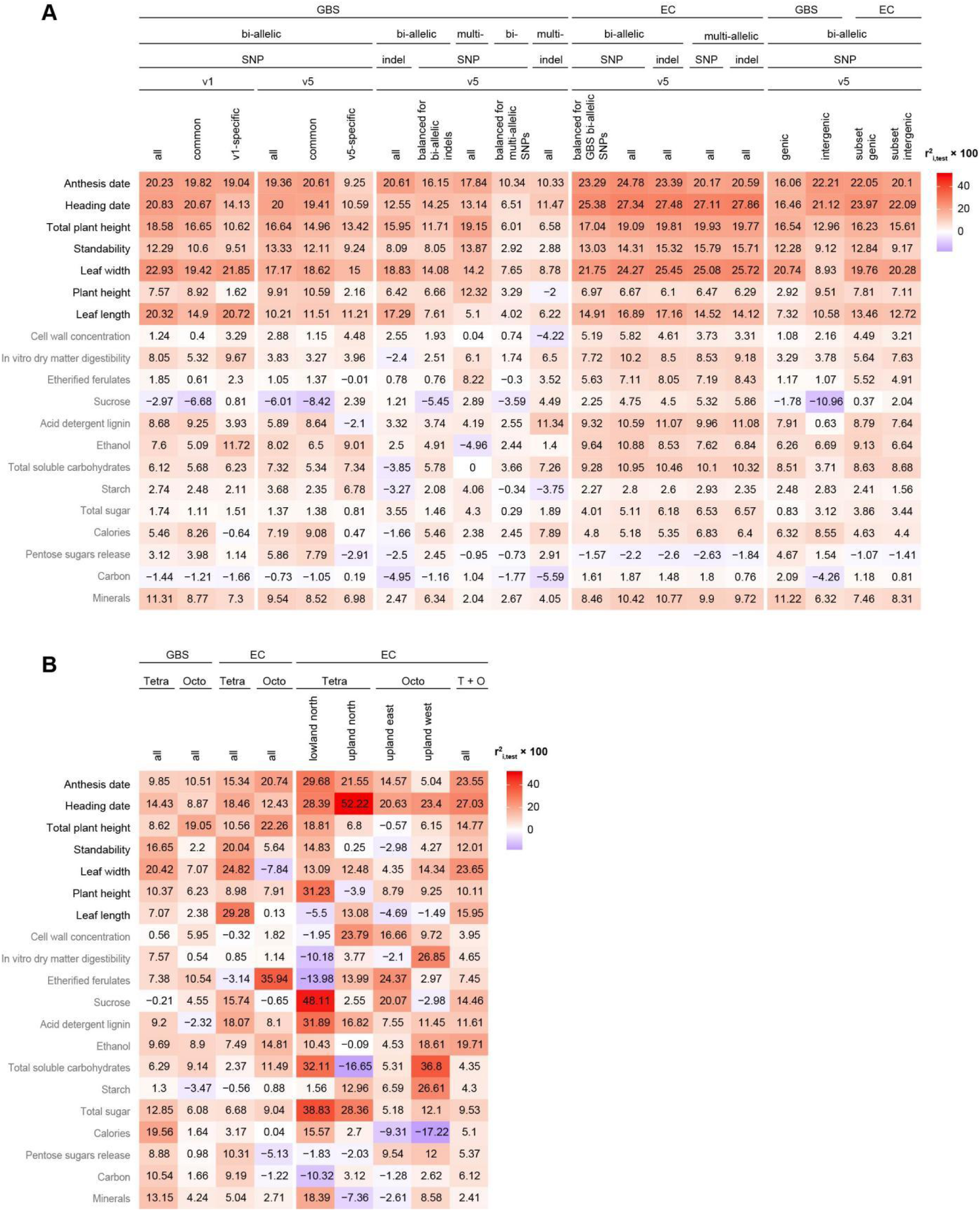
Improvement of r^2^ of models built using different genetic variants on the test sets. (**A**) r^2^ for models built on both tetraploids and octoploids using different types of variants. Table above the heatmap shows different factors the variants used may belong to. (**B**) r^2^ for models built for different ploidy levels or subpopulations using GBS or EC bi-allelic SNPs. Color scale in the heatmap: r^2^ ✕ 100.

**Fig. S8.**
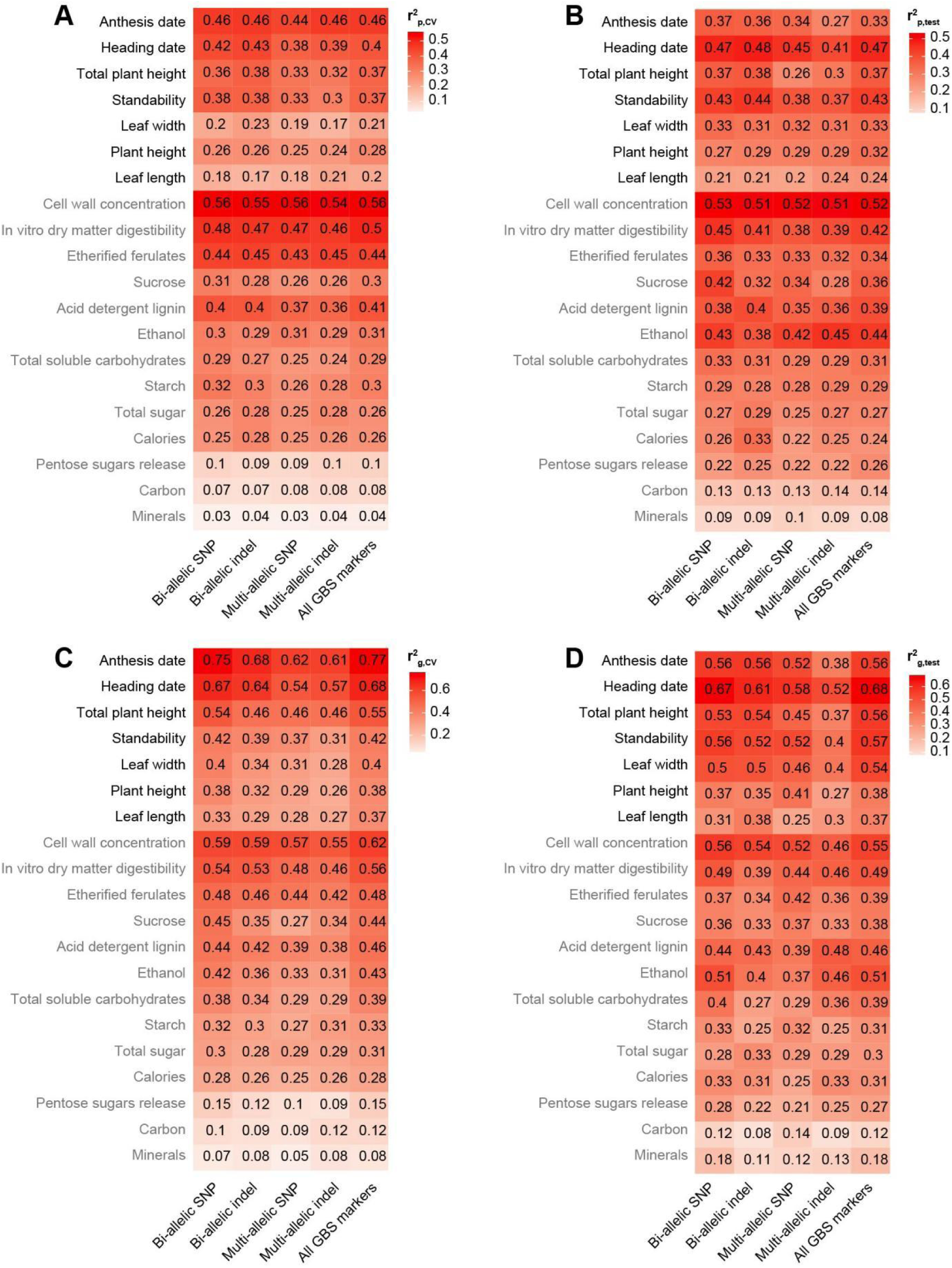
Prediction accuracy of models built using different GBS variants and models integrating all GBS variants. Color and value in (**A**-**D**) indicate r^2^_p,cv_, r^2^_p,test_, r^2^_g,cv_, and r^2^_g,test_, respectively.

**Fig. S9.**
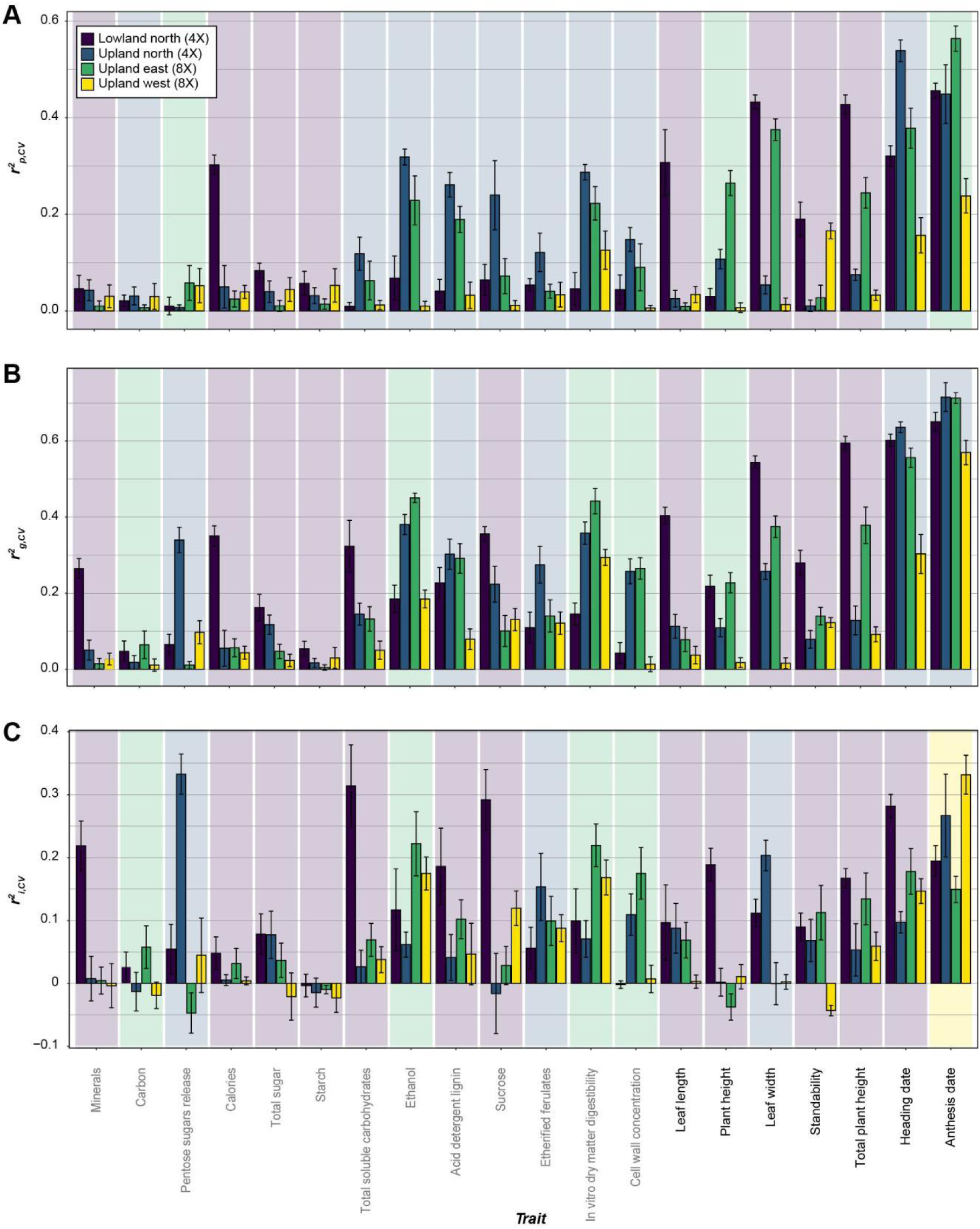
Prediction accuracy of models built for four subpopulations. (**A**-**C**) The r^2^_p,CV_ (**A**), r^2^_g,CV_ (**B**) and r^2^_i,CV_ (**C**) for models using EC bi-allelic SNPs, built for Lowland north (4X) (dark blue), Upland north (4X) (blue), Upland east (8X) (green) and Upland west (8X) (yellow) subpopulations. The color background indicates that the model built for the corresponding subpopulation had the highest prediction accuracy for the trait in question. Error bar: standard deviation.

**Fig. S10.**
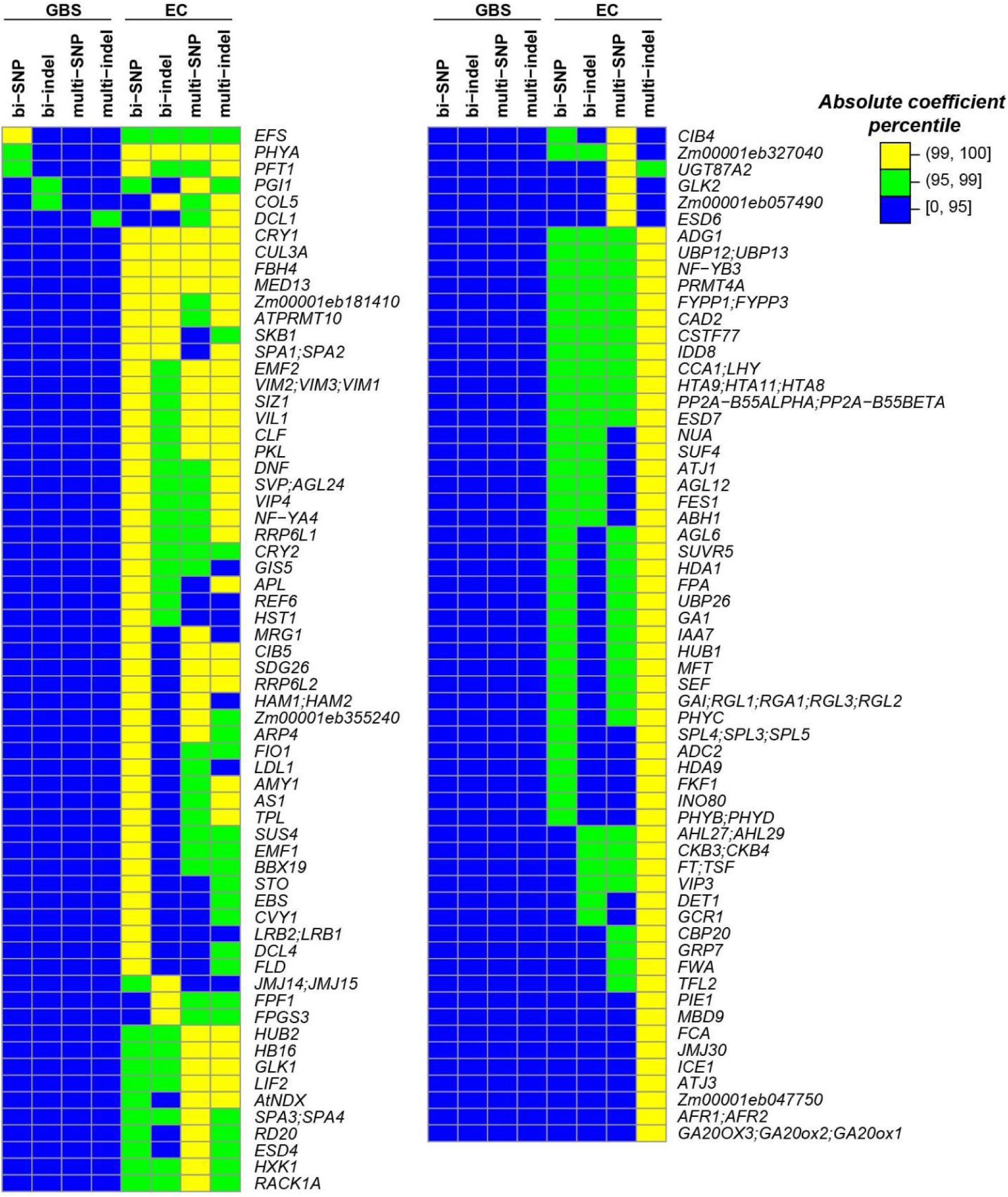
Flowering time orthologous genes associated with variants that had absolute coefficients above the 99^th^ percentile in eight models. The type of genetic variants used to build the models were listed above the heatmap. Yellow, green and blue: variants with absolute coefficients >99^th^, >95^th^ and ≤95^th^ percentiles, respectively.

